# Cells in the Polyaneuploid Cancer Cell State are Pro-Metastatic

**DOI:** 10.1101/2024.07.12.603285

**Authors:** Mikaela M. Mallin, Louis T.A. Rolle, Michael J. Schmidt, Shilpa Priyadarsini Nair, Amado J. Zurita, Peter Kuhn, James Hicks, Kenneth J. Pienta, Sarah R. Amend

## Abstract

There remains a large need for a greater understanding of the metastatic process within the prostate cancer field. Our research aims to understand the adaptive – ergo potentially metastatic – responses of cancer to changing microenvironments. Emerging evidence has implicated a role of the Polyaneuploid Cancer Cell (PACC) state in metastasis, positing the PACC state as capable of conferring metastatic competency. Mounting *in vitro* evidence supports increased metastatic potential of cells in the PACC state. Additionally, our recent retrospective study of prostate cancer patients revealed that PACC presence in the prostate at the time of radical prostatectomy was predictive of future metastatic progression. To test for a causative relationship between PACC state biology and metastasis, we leveraged a novel method designed for flow-cytometric detection of circulating tumor cells (CTCs) and disseminated tumor cells (DTCs) in subcutaneous, caudal artery, and intracardiac mouse models of metastasis. This approach provides both quantitative and qualitative information about the number and PACC-status of recovered CTCs and DTCs. Collating data from all models, we found that 74% of recovered CTCs and DTCs were in the PACC state. *In vivo* colonization assays proved PACC populations can regain proliferative capacity at metastatic sites following dormancy. Additional direct and indirect mechanistic *in vitro* analyses revealed a PACC-specific partial Epithelial-to-Mesenchymal-Transition phenotype and a pro-metastatic secretory profile, together providing preliminary evidence that PACCs are mechanistically linked to metastasis.

**Statement of Significance:** We provide the first evidence that cells in the polyaneuploid cancer cell state contribute to increased metastatic competency *in vivo*.

## I. Introduction

Though early detection of prostate cancer favors diagnosis of eradicable localized disease, metastatic prostate cancer remains lethal and incurable. Metastatic disease arises when metastatically-competent cells in the primary tumor i) invade local tissue, ii) intravasate into the vasculature, iii) survive circulatory transit, iv) extravasate into a distant organ, and v) colonize that organ (1). In 2024, it is projected that over 35,000 men in the US will die from metastatic prostate cancer (2). Clearly, there remains a large need for a greater understanding of the metastatic process within the prostate cancer field. One approach relies on understanding how tumor microenvironmental stressors constantly influence the adaptive potential of cancer cells, potentially driving phenotypes with increased metastatic potential.

Emerging evidence has highlighted the role of the Polyaneuploid Cancer Cell (or PACC) state as a phenotype of metastatically competent cells (3–7). Cells in the PACC state (also termed Polyploid Giant Cancer Cells, Multinucleate Giant Cells, and Pleomorphic Cells) exhibit a transient and adaptive cellular response to genotoxic stress. Most notably, the PACC state is characterized by an increase in genomic content coincident with an indefinite pause in cell division (8). Canonically, it is understood that genotoxic stress results in a G2/M cell cycle checkpoint stall that allows for an attempt at genomic repair, and in the event of failure, promotes apoptosis. Cells in the PACC-state adopt an interphase-restricted cell cycle following an expected stress-induced G2/M pause (9–11). This alternative cell cycle pattern, termed an endocycle, consists of subsequent cycles of G1, S, and G2 phases without any intervening M phases (12). In addition to explaining the increase in genomic content and lack of cell division, an endocycle also explains the vast nuclear and cytoplasmic enlargement typical of cells in the PACC state, which together create a distinct morphological phenotype useful in identifying PACCs in cell culture as well as histopathologic contexts. Indeed, these morphological features have been frequently used as markers of the PACC state phenotype, which has been observed to arise in various cell lines (prostate, breast, ovarian, brain, and melanoma, among others) in response to multiple classes of anti-cancer stressors (13–18).

We and others have published data supporting that cells in the PACC state have increased metastatic potential. Functionally, Xuan et. al. has reported that breast cancer MDA-MB-231 PACCs exhibit a persistent migratory phenotype driven by an enriched Vimentin filament network (19, 20). More recently, we have shown that prostate cancer PC3 PACCs demonstrate an identical motility phenotype to that observed by Xuan et al. that can be influenced by presence of a chemotactic gradient (21). In mice, Zhang et. al. showed that serial metastatic passage of PC3 cells increased not only the cells’ metastatic rates but also their percentage PACC makeup with each cycle of selection (22).

Clinical research has also indicated a potential role for PACCs in contributing to metastasis. Stromal-invasive PACCs identified by histology were more frequently identified in patients with metastatic (vs. nonmetastatic) ovarian cancer: PACCs were found in 18/21 high-grade primary tumors from patients with metastases, but only 6/26 low-grade primary tumors from patients without metastasis (23). Similar trends have been reported in prostate cancer; of 27 patients with PACC-positive cases of prostate cancer, all 27 had Gleason-scores of 9 or 10, indicating that presence of cells in the PACC state is linked to more aggressive disease (24). An independent study published nearly identical findings: of 30 patients presenting with PACC-positive cases of prostate cancer, all 30 had Gleason-scores of 9 or 10, and 11 patients were dead at a median of 8 months after diagnosis (25). The Michigan Legacy Tissue program identified PACCs in all 16 osseous and non-osseous metastatic sites of 5 randomly selected cases of prostate cancer, and most PACCs identified were concentrated around tumor hotspots (26). Most recently, we reported that presence of PACCs in the primary tumor at the time of radical prostatectomy was predictive of future metastatic progression in men with prostate cancer (27). These studies reveal a correlation between PACC state biology and metastasis.

To test for a causative relationship between PACC state biology and metastasis, we leveraged our recently published flow cytometry method designed for the detection of rare circulating tumor cells (CTCs) and disseminated tumor cells (DTCs) in metastatic mouse models (28). This approach is powerful because it provides both quantitative and qualitative information about the number and PACC-status of metastasizing CTCs/DTCs recovered from animal tissues. We used various *in vivo* models to test distinct steps of the metastatic cascade. Measurement of spontaneous metastasis of blood CTCs and distant organ DTCs from subcutaneous tumors tested invasion, intravasation, and circulation survival (for CTCs), as well as extravasation (for DTCs). Measurement of DTCs from caudal artery and tail vein injections specifically tested extravasation. Evaluation of primary tumor growth in subcutaneous models tested dormancy and colonization. Lastly, measurement of metastatic lesion outgrowth following intracardiac injection tested circulation survival, extravasation, dormancy, and colonization.

Across the various models, 75% of recovered CTCs and 72% of recovered DTCs were in the PACC state, as defined by a DNA content greater than 4N (>4N). Two *in vivo* colonization assays proved that PACC populations can regain proliferative capacity following long periods of dormancy, a phenomenon frequently observed and yet inadequately understood in the clinic. *In vitro* studies revealed a PACC-specific partial-Epithelial-to-Mesenchymal-Transition (pEMT) as a likely mechanism of increased metastatic behavior in PACCs. Notably, PACCs identified in the blood of human prostate cancer patients also demonstrated a pEMT phenotype characterized by co-expression of EpCAM and Vimentin. Additionally, an analysis of PACC-conditioned media and its effects on nonPACC cells indicated that a PACC-specific pro-metastatic secretory phenotype may increase metastatic potential of nonPACCs. Together, our results provide strong evidence that the clinically observed links between PACC presence and risk of metastasis move beyond mere correlation. Our data point to a combination of direct and indirect mechanisms that together support a causative relationship between PACCs and increased metastatic risk.

## II. Results

### The majority of circulating tumor cells are in the PACC state

We used size-matched and time-matched subcutaneous murine metastasis models to measure the differential metastatic potential of PACCs through evaluation of CTCs recovered from the blood. Mice were injected subcutaneously with either parental cells or PACC-enriched cells confirmed to have increased ploidy at the population level (Figure 1A, 2A).

**Figure 1:**
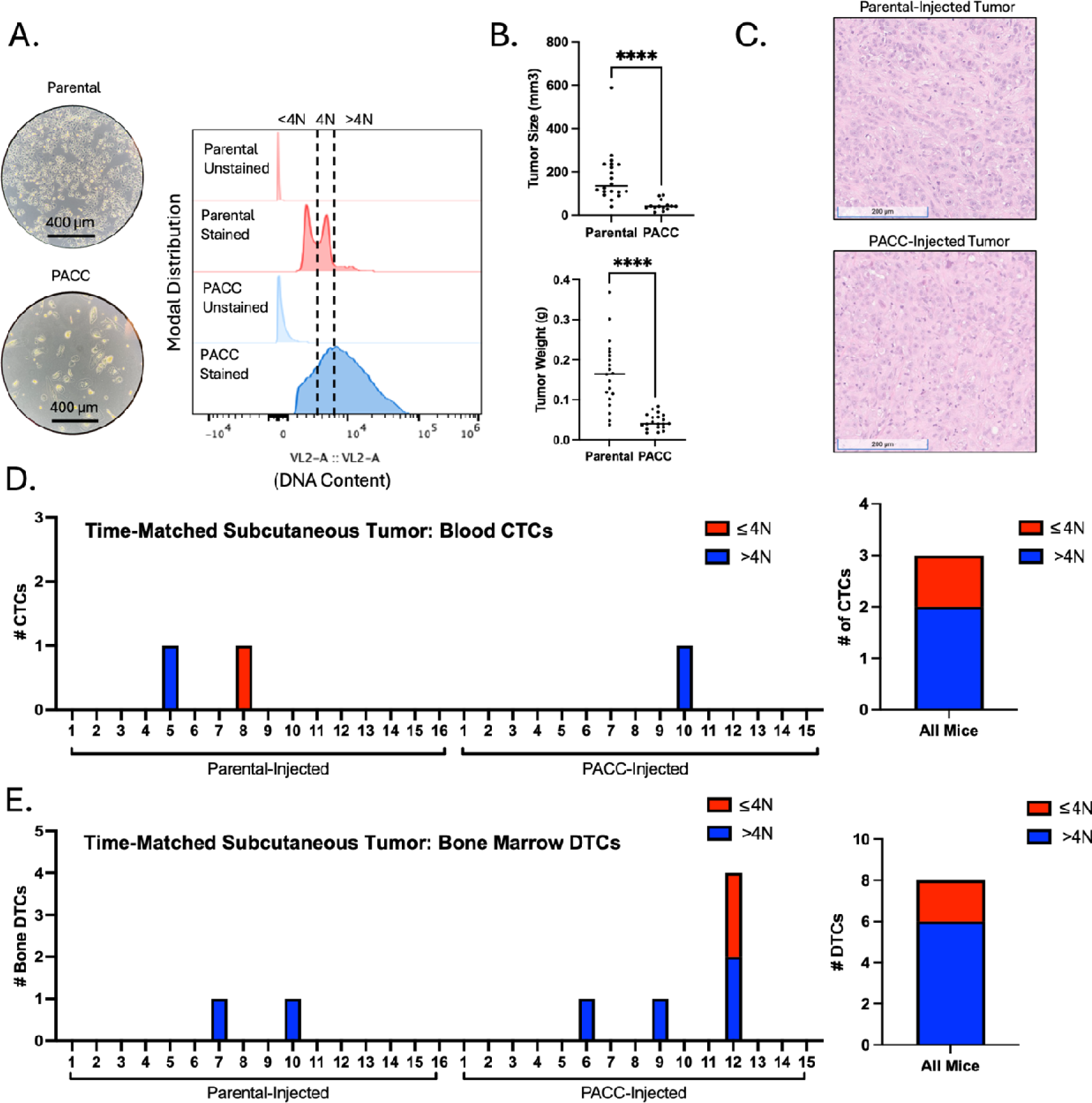
Time-matched, subcutaneous injection of PC3-GFP-Luc parental population vs. PACC-enriched population: A) Light microscopy photos and flow-cytometric ploidy analysis of injected cells per injection group. B) Tumor volume and tumor weight measurements per injection group at experimental endpoint. C) Representative H&E photos of primary tumors per injection group. D) Enumeration of CTCs sourced from the blood of each animal and quantification of % >4N CTCs vs. ≤ 4N CTCs. E) Enumeration of DTCs sourced from the bone marrow of each animal and quantification of % >4N DTCs vs. ≤ 4N DTCs.

In a time-matched model, blood from each mouse was collected and analyzed 6 weeks following tumor cell injection. At experimental endpoint, parental-injected mice produced larger tumors than PACC-injected mice (Figure 1B) probably (?) due to the transiently nonproliferative phenotype of PACCs abundant in the PACC-enriched population. At experimental endpoint, the proportion of PACCs to nonPACCs in each tumor equilibrated to similar levels (Figure 1C, appreciable by comparison of cell size). In total, 3 CTCs were recovered, 2 of which were in the PACC state (66%) (Figure 1D, Supplemental Figure S1).

To increase the number of recoverable CTCs, we repeated the experiment using a tumor size-matched model, in which the experimental endpoint of each mouse was independently determined. Blood was collected and analyzed when tumors reached approximately 350 mm^3^ (Figure 2B). Again, at experimental endpoint, the proportion of PACCs to nonPACCs in each tumor equilibrated to similar levels (Figure 2C, appreciable by comparison of cell size). In total, 33 CTCs were recovered, 25 of which were in the PACC state (75%) (Figure D, Supplemental Figure S3). Altogether, these data confirm that i) PACCs can survive as CTCs in the context of spontaneous metastasis, and ii) the majority of CTCs recovered in this context are in the PACC state.

**Figure 2:**
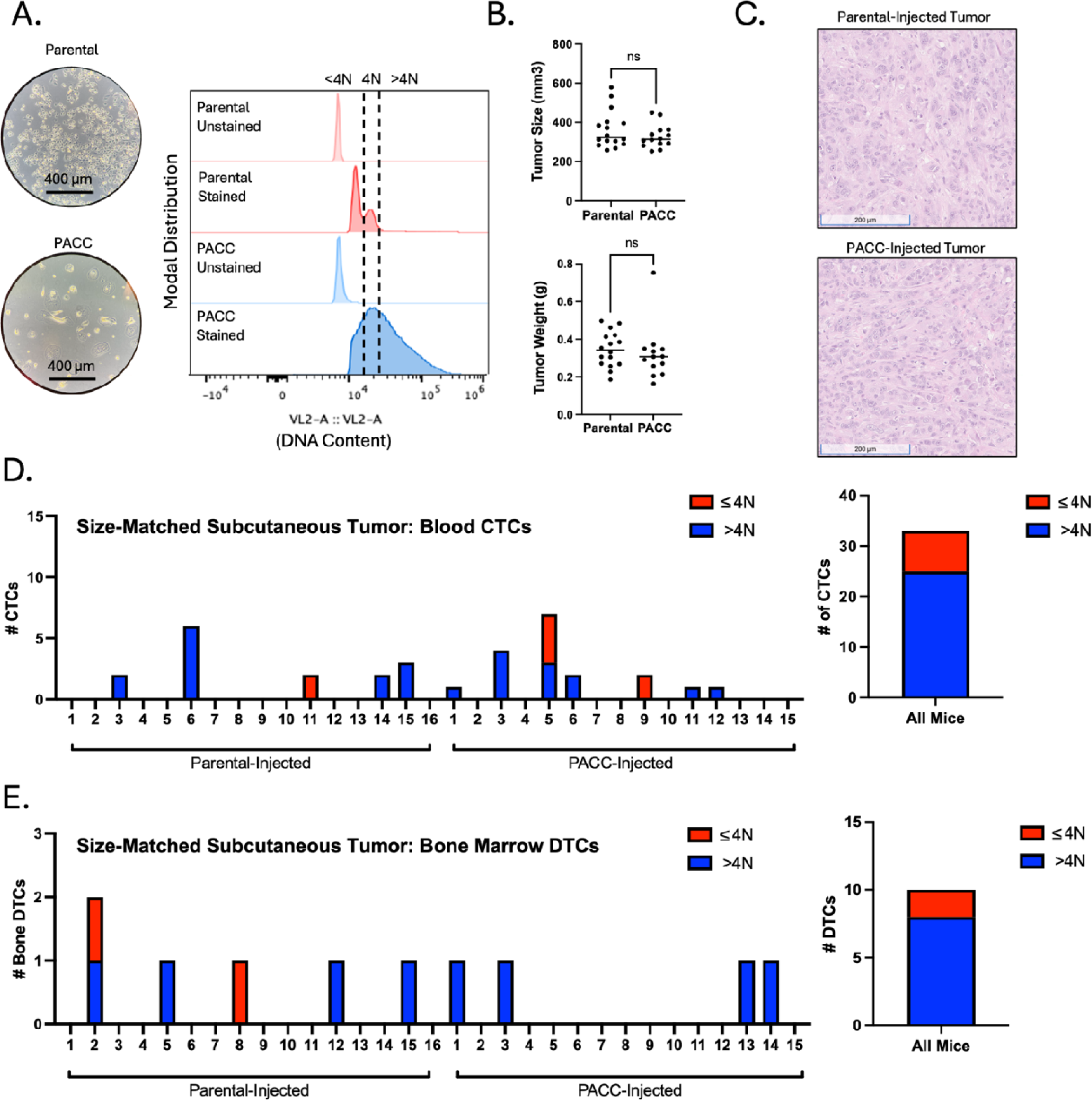
Size-matched, subcutaneous injection of PC3-GFP-Luc parental population vs. PACC-enriched population: A) Light microscopy photos and flow-cytometric ploidy analysis of injected cells per injection group. B) Tumor volume and tumor weight measurements per injection group at experimental endpoint. C) Representative H&E photos of primary tumors per injection group. D) Enumeration of CTCs sourced from the blood of each animal and quantification of % >4N CTCs vs. ≤ 4N CTCs. E) Enumeration of DTCs sourced from the bone marrow of each animal and quantification of % >4N DTCs vs. ≤ 4N DTCs.

### The majority of disseminated tumor cells are in the PACC state

The same two subcutaneous models used to measure blood CTCs were also used to quantify and characterize DTCs recovered from hind-limb bone marrow. Across both models, a majority of the DTCs recovered contained >4N DNA content, indicating they are in the PACC state. In the time-matched model, 8 bone marrow DTCs were recovered, 6 of which were in the PACC state (75%) (Figure 1E, Supplemental Figure S2). In the size-matched model, 10 DTCs were recovered, 8 of which were in the PACC state (80%) (Figure 2E, Supplemental Figure S4).

Subcutaneous tumor models are useful for investigating multiple steps of the metastatic cascade (i.e. invasion, intravasation, survival in the circulation, and extravasation) within one animal. However, they limit the ability to specifically query differential extravasation capacity between cell phenotypes owing to potential upstream bottlenecks that may differentially skew the numbers of each cell type surviving in the circulation. To directly measure the differential extravasation potential of PACCs, we used a caudal-artery injection model. Mice were injected with either parental cells or PACC-enriched cells confirmed to have increased ploidy at the population level (Figure 3A). Bone marrow and lung tissue were collected and analyzed 3 days after injection.

**Figure 3:**
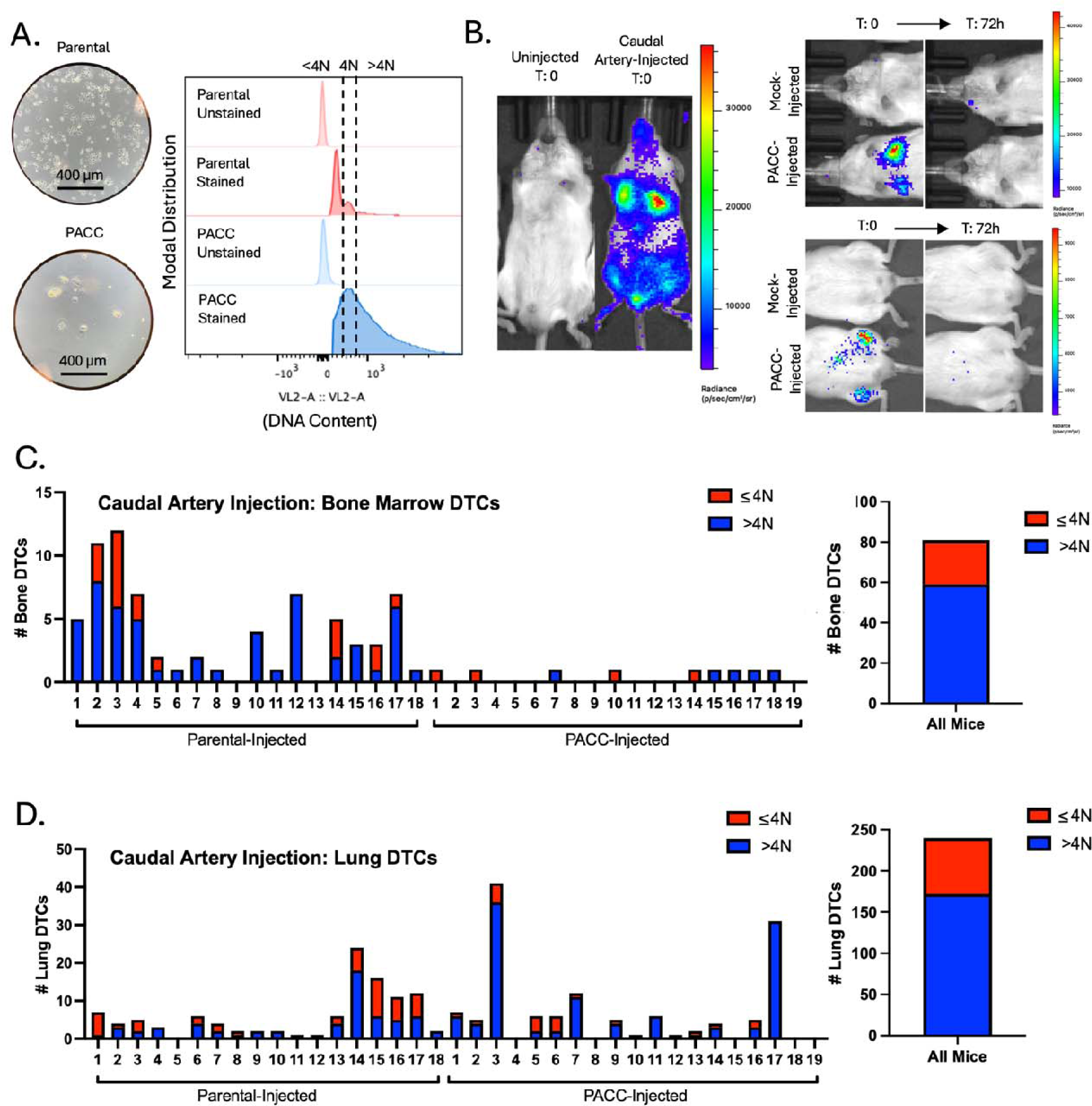
Caudal Artery injection of PC3-GFP-Luc parental population vs. PACC-enriched population: A) Light microscopy photos and flow-cytometric ploidy analysis of injected cells per injection group. B) Representative BLI images capturing the cellular distribution and signal intensity immediately following caudal artery injection and 72 hours following caudal artery injection. C) Enumeration of DTCs sourced from the bone marrow of each animal and quantification of % >4N DTCs vs. ≤ 4N DTCs. E) Enumeration of DTCs sourced from the lung tissue of each animal and quantification of % >4N DTCs vs. ≤ 4N DTCs.

Caudal artery injection introduces cells directly into the vasculature and directs them to the hind limb bone marrow capillaries and lung capillaries, wherein they become lodged due to size. After 72 hours, lodged cells have either been cleared from the vasculature or, more rarely, have extravasated into surrounding tissue (Figure 3B). After 72 hours, 81 DTCs were recovered from the bone marrow, 59 of which were in the PACC state (73%) (Figure 3C, Supplemental Figure S5). 132 DTCs were recovered from the lung, 111 of which were in the PACC state (84%) (Figure 3D, Supplemental Figure S6). This data suggests that cells in the PACC state have increased extravasation potential compared to their nonPACC counterparts. As such, not only are PACCs uniquely adept at reaching and surviving within the circulatory system as CTCs, but they are also adept at extravasating into secondary site tissues.

PACCs can be found at a low baseline level (5%) in parental populations of the PC3 cell line and are generally thought to reflect the stress inherent to cell culturing. Therefore, it is possible that PACC-state DTCs recovered from parental-injected mice result from selection of pre-existing PACCs in the parental population and reflect the increased metastatic risk of cells in the PACC state. Alternatively, it is possible that stress experienced during the metastatic process induced nonPACCs in the parental population to access the PACC state *in vivo*. To investigate the source of the PACC-state DTCs recovered from parental-injected mice, we performed a tail vein experiment comparing a population of parental cells (inherently containing a small percentage of PACCs) to a population of nonPACC enriched cells generated by size-filtering a parental population through a 10-micron filter to remove PACCs. Mice were injected with either unfiltered (parental) cells or filtered (nonPACC-enriched) cells confirmed to contain fewer cells of increased ploidy (Supplemental Figure S7A). After 72 hours, lung tissues were collected and analyzed for DTCs. Across all mice, 566 DTCs were recovered, 328 of which were in the PACC state (58%). This is consistent with our previous findings (Supplemental Figure S7B). When comparing the proportion of lung DTCs in the PACC state between the two injection groups, we found no difference (unfiltered: 58% vs. filtered: 57%) (Supplemental Figure S7C). These data show that a reduction in the percentage of baseline PACCs present in the parental population does not change the percentage of PACCs present among recovered DTCs, suggesting a possibility that nonPACCs in the parental population may access the PACC state *in vivo*.

### PACCs are capable of colonization following a period of dormancy

Clinically relevant metastatic potential requires colonization capacity. *In vitro*, cells induced to enter the PACC state become transiently nonproliferative, existing in a nondividing state. It has been observed that PACCs can survive in this state for months, a phenomenon compatible with that of metastatic tumor cell dormancy. Following a period of nonproliferative dormancy, some cells within the *in vitro* PACC-induced population return to a proliferative mitotic cell cycle, consistent with latent metastatic outgrowth frequently observed in human patients. To initially test the short-term survival status of PACCs when introduced *in vivo*, PACC-enriched cells were injected into the tail-vein of mice (Supplemental Figure S8A). The lungs were analyzed for surviving DTCs in the PACC state after 21 days (Supplemental Figure S8B). After pooling mice together, 35 DTCs in the PACC state were found still surviving in lung tissue (Supplemental Figure S8C).

Satisfied that PACCs do not immediately re-enter a proliferative cell cycle nor immediately die when introduced *in vivo*, we sought to understand the long-term survival and growth kinetics of PACCs *in vivo*. Mice were subcutaneously injected with either a population of parental cells (serving as a positive control) or a population of size filtered PACCs and monitored for 12 weeks using bioluminescent imaging (BLI). The size filtered PACC population was confirmed to contain only cells of at least 4N or greater ploidy. (Figure 4A). Within 4 weeks, 6/6 positive control mice developed appreciable tumors and showed the expected marked increase in BLI flux. At that time, 0/12 PACC-injected mice presented with palpable tumors, but BLI showed evidence of tumor cell survival at injection site (Figure 4B). Positive control mice reached ethical tumor burden and were euthanized 6 weeks post-injection. PACC-injected mice were monitored for an additional 6 weeks before they were euthanized. At experimental-endpoint, all PACC-injected mice showed greater levels of BLI flux than negative controls. 7/12 PACC-injected mice showed BLI evidence of nonproliferative tumor cell survival localized to the injection site, and an additional 2/12 PACC-injected mice had developed slow-growing palpable subcutaneous tumors following a latent dormancy phase (Figure 4C, 4D).

**Figure 4:**
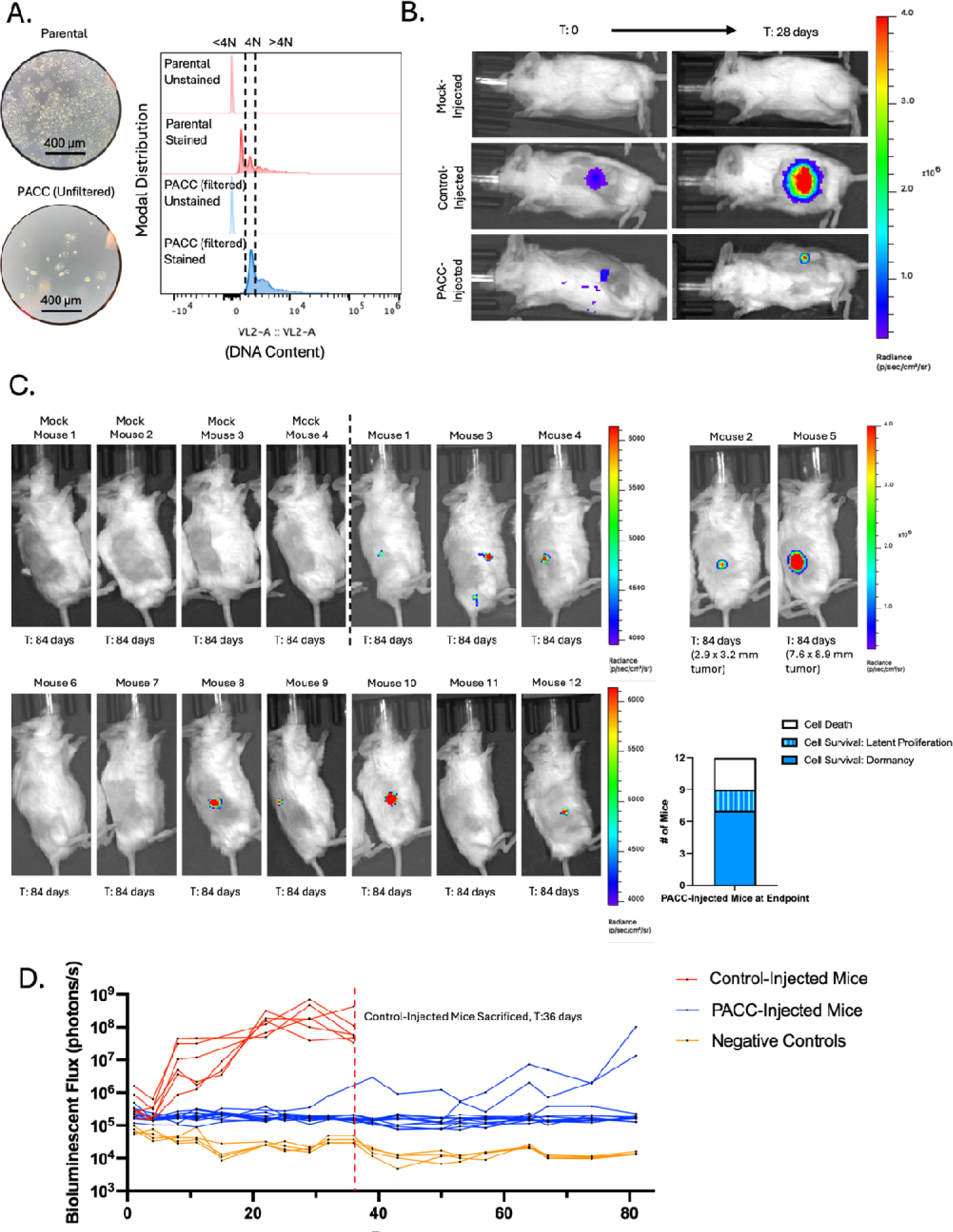
Subcutaneous injection of PC3-Luc parental population vs. size filtered PACC population: A) Light microscopy photos and flow-cytometric ploidy analysis of injected cells per injection group. B) Representative BLI images capturing the cellular distribution and signal intensity immediately following subcutaneous injection and 28 days following subcutaneous injection. C) BLI images of 4 mock-injected and 12 PACC-injected mice 82 days following subcutaneous injection. Note the different scale for Mouse 2 and Mouse 5. Quantification of the status of injected cells across all 12 PACC-injected mice at experimental endpoint. D) Weekly BLI flux of all experimental mice over 84 days.

We next used an intracardiac injection model to test the dormancy and colonization kinetics of PACCs in a metastasis-relevant context (i.e. following survival in the circulation and subsequent extravasation). Mice were injected directly into the left ventricle with either a population of parental cells (serving as a positive control) or a population of PACC-enriched cells confirmed to have increased ploidy at the population level (Figure 5A). Within 6 weeks, 6/7 positive control mice had evidence of liver and/or bone metastases by BLI. At that time, 0/7 PACC-injected mice showed any BLI positive lesions (Figure 5B). Positive control mice reached ethical tumor burden and were euthanized between 6 and 14 weeks. PACC-injected mice were monitored for an additional 5 weeks. By experimental endpoint, 4/7 PACC-injected mice had showed evidence of liver metastases by BLI, but none of these metastases reached a BLI flux threshold indicative of rapidly progressive disease (Figure 5C, 5D). Taken together, these data indicate that PACCs are capable of both i) long-term *in vivo* survival in a nonproliferative state and ii) return to a proliferative phenotype able to seed metastatic colonization following a period of dormancy.

**Figure 5:**
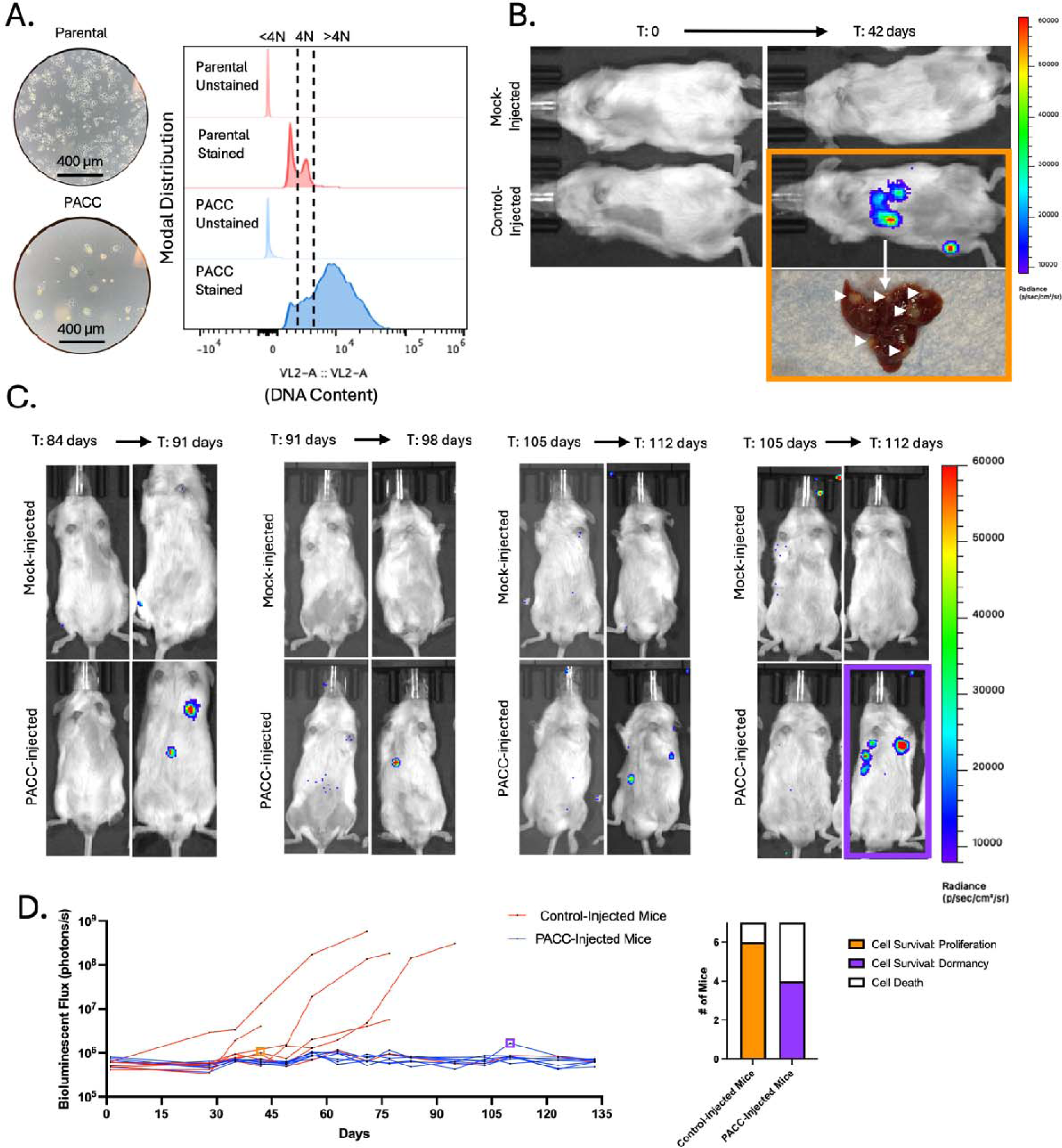
Intracardiac injection of PC3-GFP-Luc parental population vs. PACC-enriched population: A) Light microscopy photos and flow-cytometric ploidy analysis of injected cells per injection group. B) Representative BLI images capturing the cellular distribution and signal intensity immediately following intracardiac injection and 42 days following intracardiac injection. C) BLI images of 4 mock-injected and 4/7 PACC-injected mice that showed evidence of colonization following dormancy between 91- and 112-days post intracardiac injection. D) Weekly BLI flux of all experimental mice over 135 days. Quantification of the status of injected cells across all 7 PACC-injected mice at experimental endpoint.

### PACCs display a partial EMT phenotype

*In vitro* studies show that cells in the PACC state display enhanced metastatic phenotypes including motility, chemotaxis, and invasion (20, 21). The present work shows that most CTCs and DTCs recovered across various metastatic models are in the PACC state, supporting the hypothesis that PACCs promote a tumor’s metastatic potential through cell-intrinsic metastatic competency. In the past decade, it has been increasingly reported that metastatic competency relies heavily on a partial-Epithelial-to-Mesenchymal-Transition (pEMT) phenotype. pEMT (also called hybrid-EMT and mixed-EMT, among other similar names) is a nonbinarized variant of the canonically mutually exclusive epithelial vs. mesenchymal phenotypes that is characterized by co-expression of proteins typically associated with only one or the other. Using the PC3 prostate cancer cell line, we found that at the RNA level, PACCs show an increase in *ZEB1*, *VIM*, and *CLDN1* expression and no difference in *SNAI1, SNAI2, TWIST1, CDH2, CDH1,* or *EPCAM* expression (Figure 6A). At the protein level, PACCs show an increase in VIM, CDH2, CLDN1, and CDH1 expression, a decrease in SNAI1, SNAI2, and EPCAM expression, and no change in low-level TWIST1 expression (Figure 6B). High co-expression of VIM (a classic mesenchymal marker) and CLDN1 (a known epithelial marker) in PACCs indicate a pEMT phenotype. Absence of inverse expression between PACC CDH2 (a classic mesenchymal marker) and PACC CDH1 (a classic epithelial marker) also supports a PACC-specific pEMT phenotype (Figure 6C).

**Figure 6:**
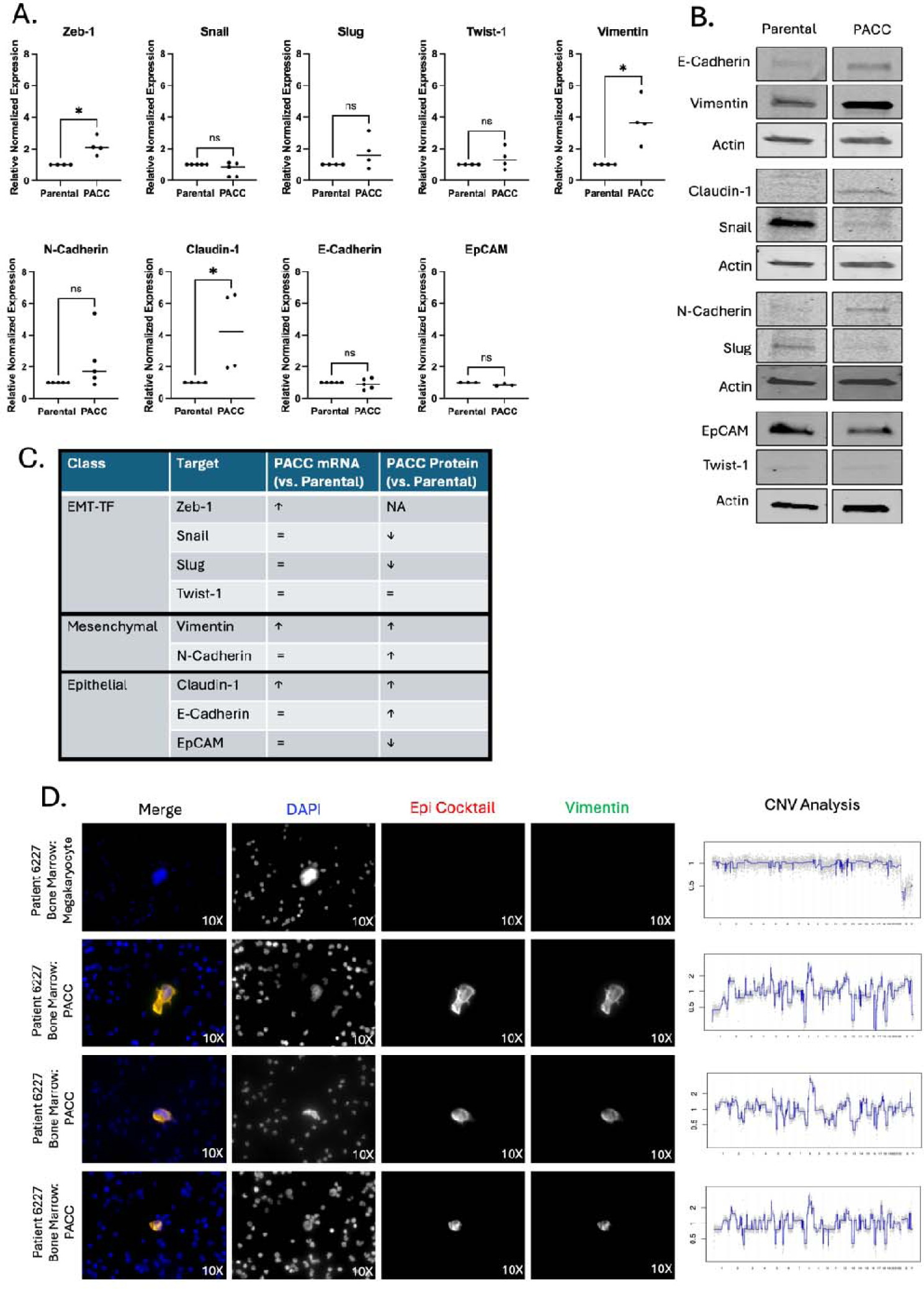
PACCs have a pEMT phenotype: A) RNA expression of a panel of EMT markers by RTqPCR in a PC3-Luc parental population vs. size filtered PACC population. B) Protein expression a panel of EMT markers by Western blot in a PC3-Luc parental population vs. size filtered PACC population, where each biological replicate reported is an average of three technical replicates. C) Summary table of RNA and protein expression. D) Representative immunofluorescent mages of PACCs identified as DTCs in the bone marrow of a metastatic prostate cancer patient, stained for DAPI, an Epithelial-origin cocktail, and VIM. CNV analysis of the corresponding single cell.

To evaluate the pEMT status of metastatic PACCs in a clinical context, we stained cancer cells in the bone marrow isolated from prostate cancer patients with a DNA content dye, a cocktail of epithelial-specific markers, and Vimentin. Cells with 4x-greater-than-average nuclear area were deemed PACCs. Single-cell copy number analysis confirmed that the PACCs were tumor-derived (Figure 6D). 34 of 44 patients were found to have PACC cancer cells. Of those 34, 6 patients were found to have pEMT PACC cancer cells positive for both pan-Epithelial (including EPCAM) and VIM stains (Representative images: Figure 6E). These data support our *in vitro* observations of PACC-specific pEMT in metastatic prostate cancer patients and provide a plausible mechanism for increased metastatic competency among cells in the PACC state.

### PACCs have a pro-metastatic secretory profile

In addition to the hypothesis that PACCs promote a tumor’s metastatic potential because PACCs themselves are more metastatic, PACCs might otherwise promote a primary tumor’s metastatic potential by increasing the metastatic phenotype of surrounding tumor cells. To test this alternative model, we evaluated the motility phenotype of nonPACC cells cultured in PACC-conditioned media compared to control parental cell-conditioned media. We found that PACC-conditioned media increases nonPACC motility by two common motility assays (Figure 7A, Supplemental Figure S9A), which was not accompanied by an increased transcriptional EMT phenotype (Supplemental Figure S9B).

**Figure 7:**
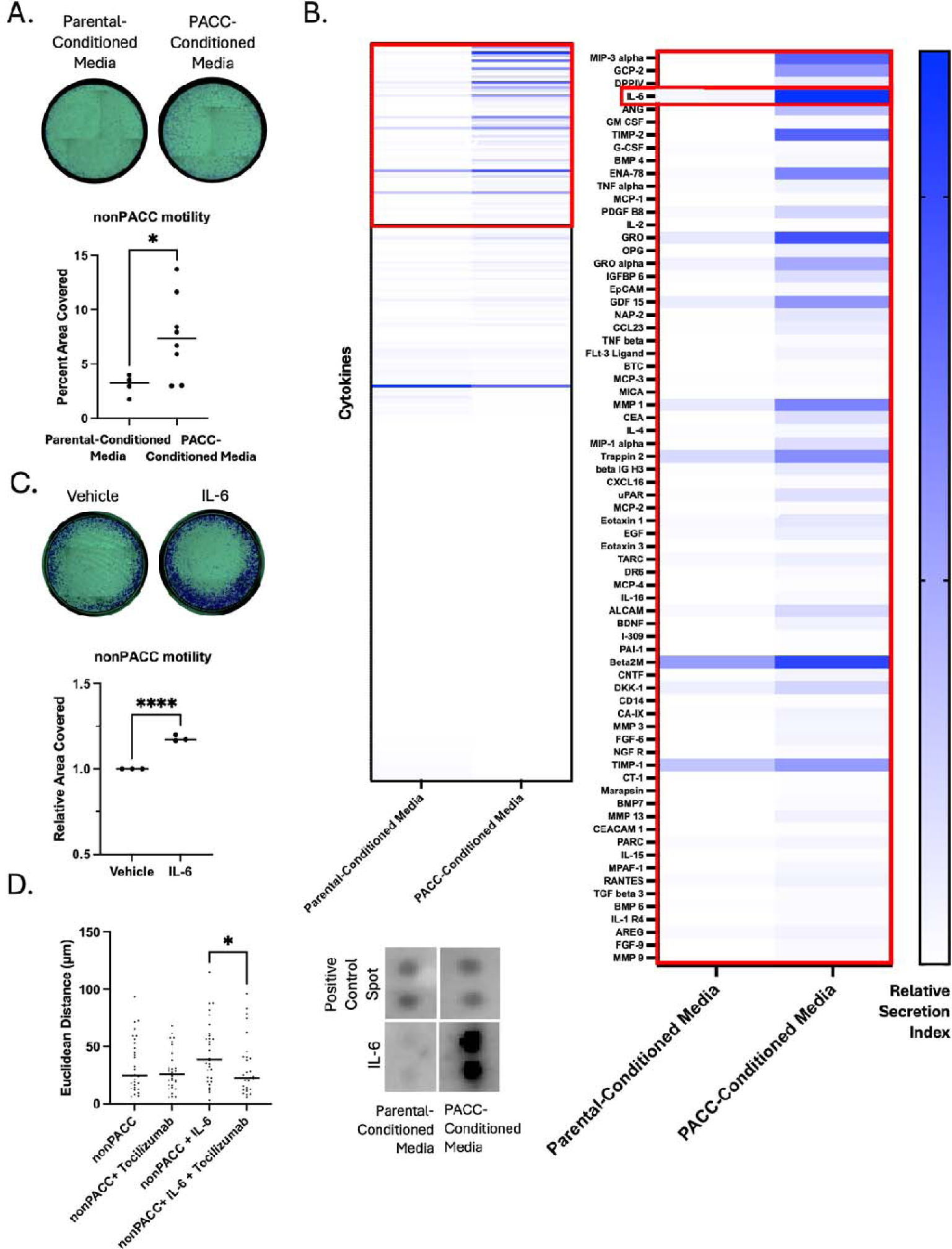
PACCs have a pro-metastatic secretory profile: A) Transwell assay to measure the differential motility of a PC3-Luc population exposed to parental-conditioned media vs. PACC-conditioned media, where each reported biological replicate is an average of two technical replicates. B) Cytokine array to measure the relative abundance of 274 cytokines of interest in parental-conditioned media vs. PACC-conditioned media. C) Transwell assay to measure the effect of addition of recombinant IL6 on the motility of a PC3-Luc population. D) Single cell tracking to measure the effects of addition of recombinant IL6 and/or Tocilizumab on the motility of a PC3-Luc population.

We performed a cytokine array on parental cell-conditioned media and PACC-conditioned media to identify abundant cytokines differentially secreted by PACCs. IL6 was the strongest candidate for further study (Figure 7B). We measured the concentration of IL6 in PACC-Conditioned Media (Supplemental Figure S9C) and found that a similar concentration was sufficient to produce increased motility in nonPACCs (Figure 7C). This increased motility phenotype was abrogated by inhibition of the IL6 Receptor (IL6R) via addition of Tocilizumab (Figure 7D). We and others have previously published the PACCs are more motile than nonPACCs (20, 21). To rule out the possibility that increased PACC motility is the result of autocrine-acting IL6, we applied Tocilizumab to PACC samples and observed no change in PACC motility (Supplemental Figure S9D). These data suggest that in addition to cell-autonomous pro-metastatic effects, PACCs contribute to a tumor microenvironment that promotes pro-metastatic phenotype in nonPACC tumor cells.

## III. Discussion

Our recent study demonstrating the clinical relevance of cells in the PACC state as reliable predictors of the risk of prostate cancer recurrence (27) raised important questions about the roles that PACCs may play in metastatic progression. For example, accession of the PACC state may increase intrinsic pro-metastatic features, making PACCs more metastatically competent than their nonPACC counterparts. Alternatively, PACCs may increase the metastatic competency of nearby nonPACCs via a paracrine-functioning pro-metastatic phenotype. Or perhaps PACCs are merely uninvolved “third-party” cells incidentally produced by an unknown stimulus that itself is driving metastatic risk via an otherwise unrelated mechanism. Here, we sought to test for a direct causative link between PACC presence and metastatic propensity. We tested for the presence of PACCs throughout various steps – and therefore locations – of the metastatic cascade using distinct metastatic mouse models.

When testing the blood for metastatic cells capable of invading, intravasating, and then surviving in the circulation, we found that 75% of recovered CTCs were in the PACC state (27/36 CTCs). When testing the bone marrow for metastatic cells capable of invading, intravasating, surviving, and then extravasating, we found that 77% of recovered DTCs were in the PACC state (14/18 DTCs). It was pertinent to test both time-matched and size-matched subcutaneous models to account for any potential effects that the differing growth kinetics of the two injections groups might have had on the initiation of the angiogenic switch. Appropriately, an increased number of total CTCs and DTCs were recovered in the size-based experiment that at endpoint had larger primary tumors than the time-based experiment. We did not consistently find that PACC-injected mice produced a larger number of metastasizing cells. Analysis of the proportion of PACCs within the tumors of each injection group at experimental endpoint offered a potential explanation: the initial difference between the proportions of PACCs within each injected population (less than 5% PACCs among parental populations, but over 50% among PACC-enriched populations) equilibrated within 6 weeks. A decrease in PACC proportion over time in the PACC-enriched tumors could be caused by a delayed return to mitotic cell cycle among PACCs, or a presence of a small number of mitotic nonPACCs in the injection population. An increase in PACC proportion over time in the parental population could be caused by accumulation of tumor microenvironmental stressors *in vivo* that initiate PACC state entry as has been previously reported (18, 29, 30).

The similarity between the proportions of PACCs found among the recovered CTCs and the recovered DTCs (75% vs 77%) in the subcutaneous tumor models does not adequately support a conclusion of increased extravasation potential among cells in the PACC state. To specifically test for increased extravasation potential among PACCs, we used a caudal artery injection model that allowed for analysis of bone marrow, the most common site of prostate metastasis in patients. When testing the bone marrow for metastatic cells capable of extravasation, we found that 73% of recovered DTCs were in the PACC state (59/81 DTCs). When testing the lung for metastatic cells capable of extravasation, we found that 84% of recovered DTCs were in the PACC state (111/132 DTCs). This data indicates that cells in the PACC state have increased extravasation potential compared to nonPACCs. We and others have previously published that PACCs demonstrate both persistent chemotactic-driven motility and functional deformability, phenotypes which may explain their increased ability to extravasate (20, 21). An alternative hypothesis supposes that extravasated nonPACCs entered the PACC state after entering secondary organ tissue owing to the inherent stressors of that organ site. When comparing the portion of PACCs found among total DTCs in the lungs of animals injected with parental populations with low-level baseline PACCs vs. a PACC-depleted population, we found that reduction of baseline PACCs present does not change the percentage of PACCs present among recovered DTCs. This data suggests that it is possible that stress experienced during the metastatic process may induce nonPACCs to access the PACC state *in vivo,* but further study is necessary.

Interestingly, in the caudal-artery model, most of the bone marrow DTCs were sourced from parental-injected mice (though the majority of those DTCs were in the PACC state), but the majority of lung DTCs were sourced from PACC-injected mice. This difference may represent biologically interesting differences in tissue-specific extravasation-barriers. For example, potential size or biophysical rigidity restrictions in the bone marrow may select for cells in the PACC state on the smaller end of what is known to be a heterogenous spectrum: the number of endocycles is inherently linked to the size of a PACC’s nucleus and cell body. Notably, baseline PACCs found in parental populations are frequently smaller than PACCs found in chemotherapy induced populations. It is possible, therefore, that bone-specific environmental pressure selects for extravasation of small PACCs, which are disproportionately found in parental populations.

Presence of CTCs and DTCs alone is not sufficient to claim complete metastatic competency; evidence of functional colonization is also needed. Multiple *in vitro* studies have reported on the latent depolyploidization (also called ploidy reversal or ploidy reduction) of cells in the PACC state following long stints of survival without cell division (31–35). The progeny of PACCs appear to be of typical cancer cell size and genomic content and display a typical mitotic cell cycle. We used two models to specifically test the dormancy and colonization potential of PACCs *in vivo.* In both settings, the PACC state life cycle closely followed what has been observed *in vitro.* 12 weeks following subcutaneous injection of size filtered PACCs (to remove any infiltrating mitotic nonPACC cells), 7/12 mice showed survival of nonproliferative PACCs at the injection site, and 2/12 mice showed delayed tumor establishment followed by slow proliferation, indicative of an *in vivo* depolyploidization event. By 19 weeks following intracardiac injection of PACCs, 4/7 mice had established, presumably proliferative, metastatic lesions.

Taken together, our *in vivo* data provides strong evidence of a causative relationship between PACCs and metastasis. Next, we sought to establish a preliminary mechanism underlying this pro-metastatic phentoype. In the past decade, there has been a notable shift away from a binary and mutually exclusive Epithelial vs. Mesenchymal phenotype. Instead, researchers have begun to appreciate the existence – and importance – of pEMT phenotypes. In fact, many groups have reported that cells with pEMT expression profiles display increased metastatic competency (36–39), perhaps owning to the needs of successfully metastatic cells to have motility programs that mesenchymal cells provide and the proliferative programs that epithelial cells provide. Our RNA and protein analysis of cells in the PACC state revealed a clear pEMT pattern, most strongly supported by simultaneous expression of *CDH1* and *VIM*. Notably, others have also reported pEMT phenotypes in PACCs (40). Analysis of bone marrow DTCs from patients with prostate cancer supported our *in vitro* findings: patient bone marrow contained PACCs co-expressing pan-Epithelial markers (including EPCAM) and VIM at the protein level.

To test for a potentially indirect mechanistic link, we turned to the literature citing the similarities between what we have termed cells in the PACC state and what others have termed therapy-induced senescent cells (41–45). The therapy-induced senescent cell literature defines this cell type as one arising in response to treatment and characterized, among other features, by a transient pause in cell cycle. Though therapy-induced senescent cells depart from classically terminal senescent cells in several ways, they have been reported to share the senescence-associated secretory phenotype (SASP). Most notably, the SASP has been reported to contribute to a more pro-metastatic tumor microenvironment (46, 47). Accordingly, we thought it possible PACCs might promote the metastatic phenotype of surrounding nonPACC tumor cells by contributing to a pro-metastatic tumor microenvironment. We found that PACCs produce a SASP-like secretory profile rich in MIP3-alpha, GCP-2, DPPIV, IL-6, GM-CSF, G-CSF, ENA-78, TNF alpha, MCP-1, IL-2, and GRO, among many others (For clarity, cytokine annotations used here are consistent with those on the product sheet). Furthermore, this co-culture with PACC conditioned media increased the motility of nonPACC cells. A validation of the most differentially secreted cytokine, IL6, showed that it was sufficient to induce motility induction in a PACC-to-nonPACC paracrine, but not a PACC-to-PACC autocrine, setting. We are not the first to find an IL6 rich PACC-secretome (48), and of note, the motility-inducing nature of IL6 has been previously published in other contexts (49).

The PACC state represents an emerging area of cancer research committed to understanding the adaptive phenotypic potential of cancer cells. It has been well established that treatment of cancer with chemotherapeutic agents inevitably leads to the eventual rise of treatment-resistant disease. Though outside the scope of this manuscript, it has been repeatedly shown that cells in the PACC state are broadly resistant to a wide swath of anti-cancer agents (8, 10, 13, 43, 48, 50, 51). In addition to chemotherapy-induced resistance, evidence has begun to suggest that cancers treated with chemotherapeutics are prone to become more metastatic. For example, emerging evidence show that chemotherapy induces cancer-cell intrinsic changes such as upregulation of anti-apoptotic genes and increased migration (52). When tested in mice using spontaneously metastatic orthotopic breast models, it was found that paclitaxel increased metastasis despite decreases in tumor burden (52). Karagiannis et. al, demonstrated that this observation holds true in the patient setting: clinically validated prognostic markers of metastasis in breast cancer patients were increased in patients who received neoadjuvant paclitaxel after doxorubicin plus cyclophosphamide (53). Considering that nearly all anti-cancer agents tested have shown to induce cells into the PACC state, our data demonstrating the increased metastatic competency of PACCs positions the PACC state as an important unforeseen ramification of neoadjuvant regimens that may help to explain clinical correlations between chemotherapy and metastatic progression.

## IV. Materials and Methodology

### Cell Culture

Experiments were performed using either the PC3-Luc prostate cancer cell line or the PC3-GFP-Luc prostate cancer cell line generated as previously described (28). All cells were cultured with RPMI 1640 media containing L-glutamine and phenol red additives (Gibco) and further supplemented with 10% Premium Grade Fetal Bovine Serum (Avantor Seradigm) and 1% 5000U/mL Penicillin-Streptomycin antibiotic (Gibco) at 37 degrees Celsius and in 5% CO2. Cells were routinely lifted using TryplE (Gibco) following a single PBS wash (Gibco). Cells were STR-profile authenticated and tested for *mycoplasma* contamination biannually (Genetica).

### PACC induction

Cells were induced to enter the PACC state as previously described (11, 21). Briefly, 625,000 PC3 cells/T75 flask (scaled for appropriately for larger tissue culture vessels) were treated with 6 µM Cisplatin (Millipore Sigma) resuspended in sterile PBS supplemented with 140mM NaCl (Sigma-Aldrich) for 72 hours. Where indicated, PC3-GFP-Luc cells were treated with 12 µM Cisplatin for 72 hours. After 72 hours, Cisplatin-treated media was removed and replaced with fresh complete media. Cells were injected (or assayed, where appropriate) 10 days after Cisplatin-treatment removal, unless indicated otherwise. When indicted, cells were filtered through a 10-micron cell strainer (PluriSelect) to either isolate or remove PACCs, which are larger in size, from the other cells in the population, as previously described (21).

### Animal Models

All murine protocols were approved by the Johns Hopkins Animal Care and Use Committee. Most experiments were performed using 8–12-week-old male Nod-Scid-Gamma (NSG) mice (Jackson Laboratories). Intracardiac injections were performed on 6-week-old mice. Subcutaneously injected mice received a 100 uL injection of a 1:1 ratio of 10 mg/mL Matrigel (Corning) to 200,000 PC3-GFP-Luc cells suspended in complete media (or a mock suspension of media alone). Caudal artery-injected mice received a 100 uL injection of 200,000 cells suspended in sterile PBS. Tail vein-injected mice received a 100 uL injection of 200,000 PC3-GFP-Luc cells suspended in sterile PBS. Intracardiac-injected mice received a 100 uL injection of 50,000 PC3-GFP-Luc cells suspended in sterile PBS into the left ventricle. Tumor progression of subcutaneously injected mice was monitored via weekly caliper measurements and tumor volume was calculated using the following formula: V = 0.5 * L * W^2^. Progression of caudal artery-injected, tail vein-injected, and intracardiac-injected mice was monitored via BLI, wherein mice were injected with 100 µL of 30 mg/mL luciferin (Regis) and imaged within 15 minutes using the IVIS Spectrum BLI imager (Revity). At experimental endpoint, subcutaneous tumors were dissection, fixed in 10% Neutral Buffered Formalin (Sigma-Aldrich) for 24 hours, washed 3x 5 minutes in PBS, and embedded into paraffin blocks. 4-micron thick sections were mounted on slides, stained with Hematoxylin and Eosin, and imaged using a 40X objective.

### CTC and DTC Analysis

CTC and DTC detection and analysis was performed as previously described (28). CTCs were defined as GFP-positive cells identified in mouse blood by flow cytometry. 500 µL of blood was collected via terminal tail bleed. Blood samples were individually transferred to 5 mL Eppendorf tubes and supplemented with ACK lysis buffer (Quality Biological) at a 1:4 blood:lysis buffer ratio. This solution was incubated on an end-over-end turner for 10 minutes. Following incubation, all samples were centrifuged at 1500 xg for 10 minutes at 4 degrees C. Cell pellets were resuspended in 1 mL complete media and stained with 1 µL of Vybrant DyeCycle Violet (Thermo Fisher Scientific).

DTCs were defined as GFP-positive cells identified in mouse hind-limb blood marrow or homogenized lung tissue by flow cytometry. Bone marrow was collected from the hind limb bones of freshly euthanized mice using a standard centrifugation protocol. Briefly, the right and left femurs and tibias of each mouse were dissected. The distal femoral epiphysial plate from each femur and the proximal tibial epiphysial plate from each tibia were removed to ensure access to red bone marrow. Bones were placed marrow-exposed side down in a 0.5 mL tube punctured with a small hole which was then nested into a 1.5 mL tube. Tubes were centrifuged at maximum speed for 30 seconds to collect bone marrow into the 1.5 mL tube. All samples were resuspended in 200 μL PBS and then supplemented with 800 μL of ACK lysis buffer. This solution was incubated on an end-over-end turner for 10 minutes. Following incubation, all samples were centrifuged at 1500 xg for 10 minutes at 4 degrees C. Cell pellets were resuspended in 3 mL complete media and stained with 15 µL of Vybrant DyeCycle Violet.

To collect lung tissue, all 5 lobes of the lungs were dissected from freshly euthanized mice. Each sample was transferred to a petri dish and minced into small pieces with a fresh, straight-edged razor blade. Each sample was transferred to a 14 mL round-bottom tube and suspended in 5 mL of a solution containing 250 µL Collagenase/Hyaluronidase (Stem Cell Technologies), 375 µL DNase I (Stem Cell Technologies) at 1 mg/mL, and 1.875 mL complete media. The samples were incubated at 37 degrees C for 20 minutes with shaking, before being pushed through a fresh 70-micron strainer placed over a 50 mL conical tube using the rubber end of a fresh 5 mL syringe plunger. 45 mL of complete media was used per sample to facilitate straining. Following straining, samples were centrifuged at 300 xg for 10 minutes at room temperature. The cell pellets were resuspended in 2 mL of ACK lysis buffer and incubated at room temperature for 3 minutes before adding 47 mL of complete media. Samples were then centrifuged at 300 xg for 10 minutes at room temperature, using a slow deceleration setting. Cell pellets were resuspended in 2 mL of complete media and stained with 10 µL of Vybrant DyeCycle Violet.

The entire volume of all stained samples was run on the Attune NxT Acoustic Focusing Cytometer (Thermo Fisher Scientific) at a flow rate of 1000 µL per minute. As previously described, it was critical to implement the following four modifications to our cytometer to accommodate analysis of cells in the PACC state: 1) The largest commercially available blocker bar was installed. 2) An alternative optical configuration was used. 3). Thresholding was performed using SSC rather than the standard FSC. 4) Area scaling factors of 0.6 were used for all lasers, rather than the standard area scaling factors. With these modifications, data was collected in the SSC, VL1-A, VL2-A, and BL1-A channels. All data were analyzed using FlowJo (BD), following our previously published analysis protocol (28).

### Patient bone marrow samples

Bone marrow aspirate (BM) samples were collected from castrate-resistant prostate cancer patients as previously described (54). Briefly, liquid biopsy samples were taken from participants at baseline and immediately started treatment on trial NCT01505868 which evaluated cabazitaxel with or without carboplatin. Samples were collected at MD Anderson at baseline prior to clinical trial treatment administration. The study was approved by the corresponding institutional review boards and was conducted in accordance with ethical principles founded in the Declaration of Helsinki. All patients gave written informed consent.

### Patient sample processing

Samples were processed as previously described (54). In short, bone marrow samples were delivered overnight (7.5 mL) in Cell-Free DNA Blood Collection Tubing (Streck) at the clinical sites and sent to the University of Southern California for processing. Erythrocytes were lysed via ammonium chloride and the entire nucleated cell population was plated onto a specialized cell adhesion glass slide (Marienfield) at a density of 2-3 million cells per slide. Slides were stored at −80°C until use.

### Immunofluorescent staining

Slides were fixed with 2% PFA for 20 minutes. Slides were then blocked with 2% BSA in PBS. Slides were then incubated overnight at 4°C with an antibody cocktail consisting of mouse IgG1/Ig2a anti-human cytokeratins (CK) 1, 4, 5, 6, 8, 10, 13, 18, and 19 (clones: C-11, PCK-26, CY-90, KS-1A3, M20, A53-B/A2, C2562, Sigma, St. Louis, MO, USA), mouse IgG1 anti-human CK 19 (clone: RCK108, GA61561-2, Dako, Carpinteria, CA, USA), mouse EpCAM (14-9326-82, Thermo), and rabbit IgG antihuman vimentin (VIM): Alexa Fluor 488 (clone: D21H3, 9854BC, Cell Signaling Technology). Slides were then washed with PBS and incubated at room temperature for two hours with Alexa Fluor® 555 goat anti-mouse IgG1 antibody (A21127, Invitrogen, Carlsbad, CA, USA), and counter-stained with 4′,6-diamidino-2-phenylindole (DAPI; D1306, Thermo Fisher Scientific, Waltham, MA, USA).

### High content imaging and analysis

Slides were imaged as previously reported (55). Briefly, an automated high throughput microscope equipped with a 10x optical lens was used to collect 2304 images across the slide. An image analysis tool, available at https://github.com/aminnaghdloo/if_utils was used to identify PACCs. Briefly, each fluorescent channel was segmented individually using adaptive thresholding and merged into one cell mask. PACCs were identified as having a nuclear diameter two-times larger than the rare cell population (four-times the area).

### Single cell picking

AN Eppendorf TransferMan NK2 micromanipulator was used to collect the cell of interest in a 100 µM micropipette. The cell was transferred into a sterile solution of 1x PBS. Single cells were stored at −80°C.

### Copy number profiling

Copy number profiling from low pass whole genome sequencing samples was conducted as previously described (56, 57). Briefly, the commercially available WGA4 kit (Sigma) was used for single cell whole genome amplification. Sequencing libraries were prepared with the NEB Ultra FS II with 50 ng of starting material. Cells were sequenced at a depth of 1-2 million reads on an Illumina HiSeq 4000 (Fulgent, Inc.). Raw sequencing reads were aligned with BWA-MEM to the hg19 reference. Count data was segmented via the R package DNACopy (version 1.70.0) and median values were reported for copy number ratio data.

### Western Blot

PC3 parental cells or PC3 PACCs were lysed with an appropriate amount of RIPA Lysis and Extraction Buffer (Thermo Scientific) with Halt Protease and Phosphatase Inhibitor Cocktail (Thermo Scientific) for 30 minutes, rotating in 4 degrees Celsius. Lysates were spun at 21,000 xg for 15 minutes in 4 degrees Celsius and the proteinaceous supernatant was stored at −80 degrees Celsius. 50 ng of protein (measured by Pierce BCA Protein Assay, following manufacturer’s protocol) (Thermo Scientific) was added to a 1:4 mixture of Laemmli Sample Buffer (BioRad) and 2-Mercaptoethanol (BioRad) and ran through a 4-20% Mini-ProTEAN TGX gel (BioRad). The gel was transferred via Trans-Blot SD Semi-Dry Transfer Cell (BioRad) onto a 0.2-micron Nitrocellulose Trans-Blot Turbo Transfer Pack using the 7-minute, 2.5A, 25V protocol designed for Mixed Molecular Weights. The blot was blocked in Casein Blocking Buffer (Sigma-Aldrich) for 1 hour at room temperature with shaking, and then transferred to primary antibody diluted in casein and incubated overnight at 4 degrees Celsius with shaking. The blot was then washed 3 times for 5 minutes each with pH 7.4 Tris-Buffered Saline (Quality Biological) with 0.1% Tween 20 (Sigma) (TBST) and incubated in secondary antibody diluted 1:20,000 in Casein for 1 hour at room temperature. The blot was then washed 3 times for 5 minutes with TBST and imaged using the Odyssey Western Blot Imager (Li-Cor). See Table 1 below for antibodies used.

**Table 1:** Western Blot Antibodies.

| **Target**<br>**(Alias)** | **Antibody** | **Concentration** | **Secondary** |
| --- | --- | --- | --- |
| SNAI1<br>(Snail) | Rabbit IgG Monoclonal: C15D3<br>(Cell Signaling Technologies 3879) | 1:1000 | IRDye 800CW Goat anti<br>Rabbit IgG (Licor) |
| SNAI2<br>(Slug) | Rabbit IgG Monoclonal: C19G7<br>(Cell Signaling Technologies 9585) | 1:1000 | IRDye 800CW Goat anti<br>Rabbit IgG (Licor) |
| TWIST1<br>(Twist-1) | Rabbit IgG Monoclonal: E7E2G<br>(Cell Signaling Technologies 31174) | 1:500 | IRDye 800CW Goat anti<br>Rabbit IgG (Licor) |
| VIM<br>(Vimentin) | Rabbit IgG Monoclonal: D21H3<br>(Cell Signaling Technologies 5741) | 1:5000 | IRDye 800CW Goat anti<br>Rabbit IgG (Licor) |
| CDH2<br>(N-Cadherin) | Rabbit IgG Monoclonal: D4R1H<br>(Cell Signaling Technologies 13116) | 1:500 | IRDye 800CW Goat anti<br>Rabbit IgG (Licor) |
| CLDN1<br>(Claudin-1) | Rabbit IgG Monoclonal: D5H1D<br>(Cell Signaling Technologies 13255) | 1:1000 | IRDye 800CW Goat anti<br>Rabbit IgG (Licor) |
| CDH1<br>(E-Cadherin) | Rabbit IgG Monoclonal: 24E10<br>(Cell Signaling Technologies 3195) | 1:1000 | IRDye 800CW Goat anti<br>Rabbit IgG (Licor) |
| EPCAM<br>(EpCAM) | Rabbit IgG Monoclonal: E678Y<br>(Cell Signaling Technologies 93790) | 1:1000 | IRDye 800CW Goat anti<br>Rabbit IgG (Licor) |
| ACTB<br>(Beta-Actin) | Mouse IgG2B Monoclonal:<br>8H10D10 (Cell Signaling<br>Technologies 3700) | 1:5000 | IRDye 680RD Goat anti<br>Mouse IgG (Licor) |

### RT-qPCR

PC3 parental cells or PC3 PACCs were lysed using the QIAshredder Kit (Qiagen), following the manufacturer’s protocol. Where indicated, PC3 parental cells or PC3 PACCs were incubated in parental-conditioned media or PACC-conditioned media for 24 hours prior to lysis. RNA was extracted from lysates using an RNeasy Mini Kit (Qiagen) following the manufacturer’s protocol. RNA was converted to cDNA (1 ug RNA per reaction) using the iScript cDNA Synthesis Kit (Bio-Rad) following the manufacturer’s protocol. RT-qPCR reactions were performed using SsoFast EvaGreen Supermix (Bio-Rad), following the manufacturer’s protocols, and assorted primers (Integrated DNA Technologies) at a concentration of 10 uM. See Table 2 below for primers used. Data was collected using the CFX96 Real-Time PCR Detection System (Bio-Rad) with a standard cycle protocol. Gene expression was normalized to the housekeeping gene Beta-Actin and calculated using the delta-delta Ct method. Each biological replicate reported is an average of three technical replicates.

**Table 2:**
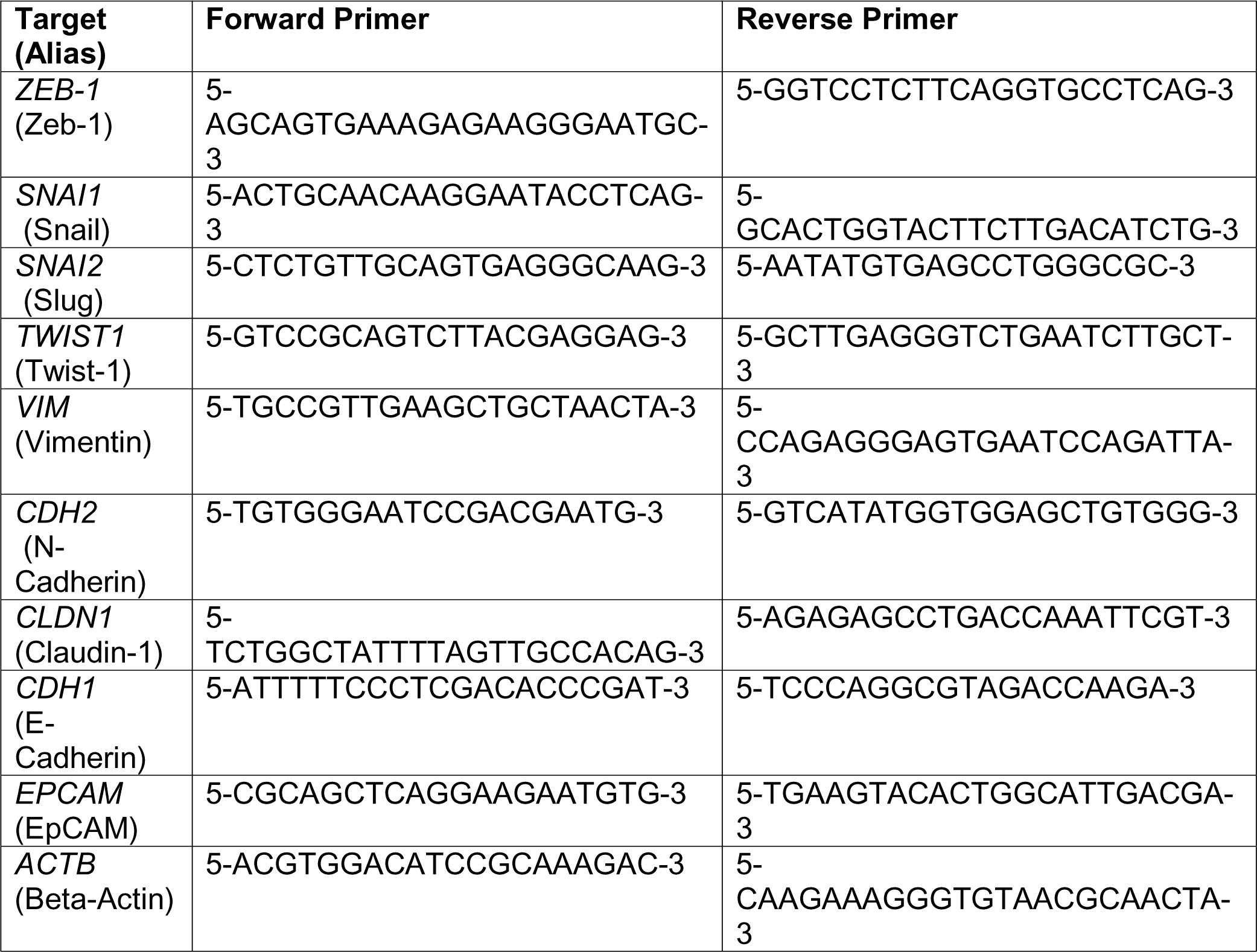
RT-qPCR Primers.

### Conditioned-Media Generation

To generate PACC-conditioned media, PC3 PACCs were generated using our standard induction-approach as described above, in a T150 tissue-culture flask. On day 10 of treatment-removal, exactly 20 mL of fresh complete media were added. Exactly 24 hours later, on day 11 of treatment-removal, all 20 mL of PACC-conditioned media was collected. Media was centrifuged at 1000 xg for 5 minutes to remove any debris, and then filtered through a 0.45-micron PES filter into 1 mL aliquots (to prevent freeze-thawing) that were stored at −80 degrees Celsius.

To generate parental-conditioned media, 1,250,000 PC3 parental cells were seeded in a T150 tissue-culture flask. After 12 hours, exactly 20 mL of fresh complete media were added. Exactly 24 hours later, all 20 mL of parental-conditioned media was collected. Media was centrifuged at 1000 xg for 5 minutes to remove any debris, and then filtered through a 0.45-micron PES filter into 1 mL aliquots (to prevent freeze-thawing) that were stored at −80 degrees Celsius.

### Motility Assays

Transwell assays were performed using 6.5-millimeter, 8-micron, PES membranes. PC3 parental cells were seeded on the membrane at 100,000 cells/mL in 250 µL of complete media. After 12 hours, 500 µL of testing-condition media was placed in the top and bottom wells for 24 hours. Where indicated, recombinant IL6 was added to complete media at a concentration of 0.5 ng/mL. After 24 hours, membranes were washed in PBS and nonmigrated cells were removed from the top of the membrane with a cotton-tipped applicator. Membranes were fixed in 100% ice cold methanol for 10 minutes, and then stained in 0.5% crystal violet resuspended in 20% methanol for 10 minutes. Membranes were washed with deionized water to remove excess stain and tiled imaging of the entire membrane was acquired with an EVOS FL Digital Inverted Fluorescence Microscope (Thermo Fisher Scientific) using a 4X objective. Image analysis to calculate percent area coverage was performed using the Phansalkar auto-local threshold method in ImageJ. Each reported biological replicate is an average of two technical replicates.

Wound healing assays were performed in 24-well tissue culture plates. PC3 parental cells were seeded at 100,000 cells/mL in 2.5 mL of complete media. After 12 hours, a P1000 pipette tip was used to create a scratch wound. After 3 washes with PBS, the scratch wounds were imaged on an EVOS FL Digital Inverted Fluorescence Microscope using a 4X phase objective. 1mL of testing-conditioned media was added for 24 hours, after which the wounds were re-imaged. Image analysis to calculate percent wound closure was performed using the Wound Healing Size Tool Updated plug-in in ImageJ. Each reported biological replicate is an average of two technical replicates.

Single-cell tracking assays were performed using live-cell, time-lapse microscopy in 24-well tissue culture plates. PC3 parental cells were seeded at 25,000 cells/mL in 2 mL of complete media. PC3 PACCs were seeded 5,000 cells/mL in 2 mL of complete media. After 12 hours, testing-condition media was added for 24 hours. Where indicated, Tocilizumab (Selleck Chemicals) was added to complete media at a concentration of 2.5 µg/mL. Where indicated, Recombinant IL6 was added to complete media at a concentration of 0.5 ng/mL. An EVOS FL Digital Inverted Fluorescence Microscope was used to take 10X phase images every 30 minutes for the 24-hour testing-condition incubation. An on-stage environment chamber was used to maintain cell conditions at 37 degrees Celsius, 5% CO_2_, and 20% O_2_. Images were analyzed using the Manual Tracking and Chemotaxis Tool plug-ins in ImageJ image analysis software. All cells analyzed were randomly selected. Cells that underwent division, apoptosis, or moved out of frame were excluded from analysis.

### Cytokine Array

Cytokine analysis was performed using a 274-target chemiluminescent human cytokine antibody array, following manufacturer’s protocol. (Abcam ab198496). Nondiluted PACC-conditioned media and Parental-conditioned media were tested, using nonconditioned media as a background control. Membranes were imaged using the ChemiDoc XRS+ imaging system (Biorad) and densitometry data was obtained using Image Lab Software (Biorad) with signal-specific automatic background thresholding enabled. Background subtraction, positive control normalization differential secretion calculations were performed following manufacturer’s protocol.

### Elisa

Quantification of IL6 present in PACC-Conditioned media was performed using an IL6 Elisa, following the manufacturer’s protocol (BioLegend 430504), with the following alteration: sample incubation time was lengthened from 2 hours to 3 hours. PACC-Conditioned media was diluted 1:4 in protocol buffer Assay A prior to analysis. Data was collected using FLUOStar Omega Plate Reader (BMG LabTech) at 405 nm.

### Statistics

Nonparametric T-Tests (Mann-Whitney) were performed to generate reported P values. Power calculations were performed to determine appropriate sample size for *in vivo* experiments, wherein an alpha value of 0.05 and a beta value of 0.8 were used.

NS = nonsignificant = P > 0.05.

* = P < 0.05

** = P < 0.01

*** = P < 0.001

**** = P < 0.0001

## Authors’ Contributions

M.M.M: Conceptualization, Formal Analysis, Investigation, Methodology, Validation, Visualization, Writing – Original Draft; L.T.A.R: Investigation, Writing – Review and Editing; M.J.S: Formal Analysis, Investigation, Visualization, Writing – Review and Editing; S.P.N: Investigation, A.J.Z: Writing-Review and Editing; P.K: Funding Acquisition, Writing – Review and Editing; J.H: Supervision, Funding Acquisition; K.J.P: Conceptualization, Funding Acquisition, Writing – Review and Editing; S.R.A: Funding Acquisition, Writing – Review and Editing.

## Competing Interests

M.M.M. has no disclosures. L.T.A.R. has no disclosures. M.J.S has no disclosures. S.P.N. has no disclosures. ruA. J. Z. has no disclosures. P.K. discloses ownership in Epic Sciences. J.H discloses he is a member of the Clinical Advisory Board of Epic Sciences. K.J.P. discloses that he is a consultant to Cue Biopharma, Inc., an equity holder in PEEL therapeutics, and a founder and equity holder in Keystone Biopharma, Inc. and Kreftect, Inc. S.R.A. discloses that she is an equity holder in Keystone Biopharma, Inc.

## Ethics Statement

All animal experiments were reviewed and approved by the Johns Hopkins Animal Care and Use Committee. Human elements of this study were approved by the corresponding institutional review boards and were conducted in accordance with ethical principles founded in the Declaration of Helsinki. All patients gave written informed consent.

## Funding

P.K and J.H were supported by the National Cancer Institute’s Norris Comprehensive Cancer Center (CORE) Support 5P30CA014089-40. K.J.P was supported by National Cancer Institute grants U54CA143803, CA163124, CA093900, and CA143055, and the Prostate Cancer Foundation. S.R.A. was supported by the US Department of Defense CDMRP/PCRP (W81XWH-20-10353 and W81XWH-22-1-0680), the Patrick C. Walsh Prostate Cancer Research Fund, and the Prostate Cancer Foundation.

## Acknowledgements

The authors would like to acknowledge Ana Aparicio and Paul Corn for their critical contributions. The authors would also like to thank the patients and their caregivers who consented to this study, as well as the clinical research staff who contributed. The authors also thank the members of the Cancer Ecology Center for thoughtful conversation and invaluable feedback.

## Data Availability Statement

Nearly all data generated in this study are available within the article and its supplementary data files. Detailed data regarding the 247-panel cytokine array are available upon request from the corresponding author.

---

**Supplemental Figure S1:**
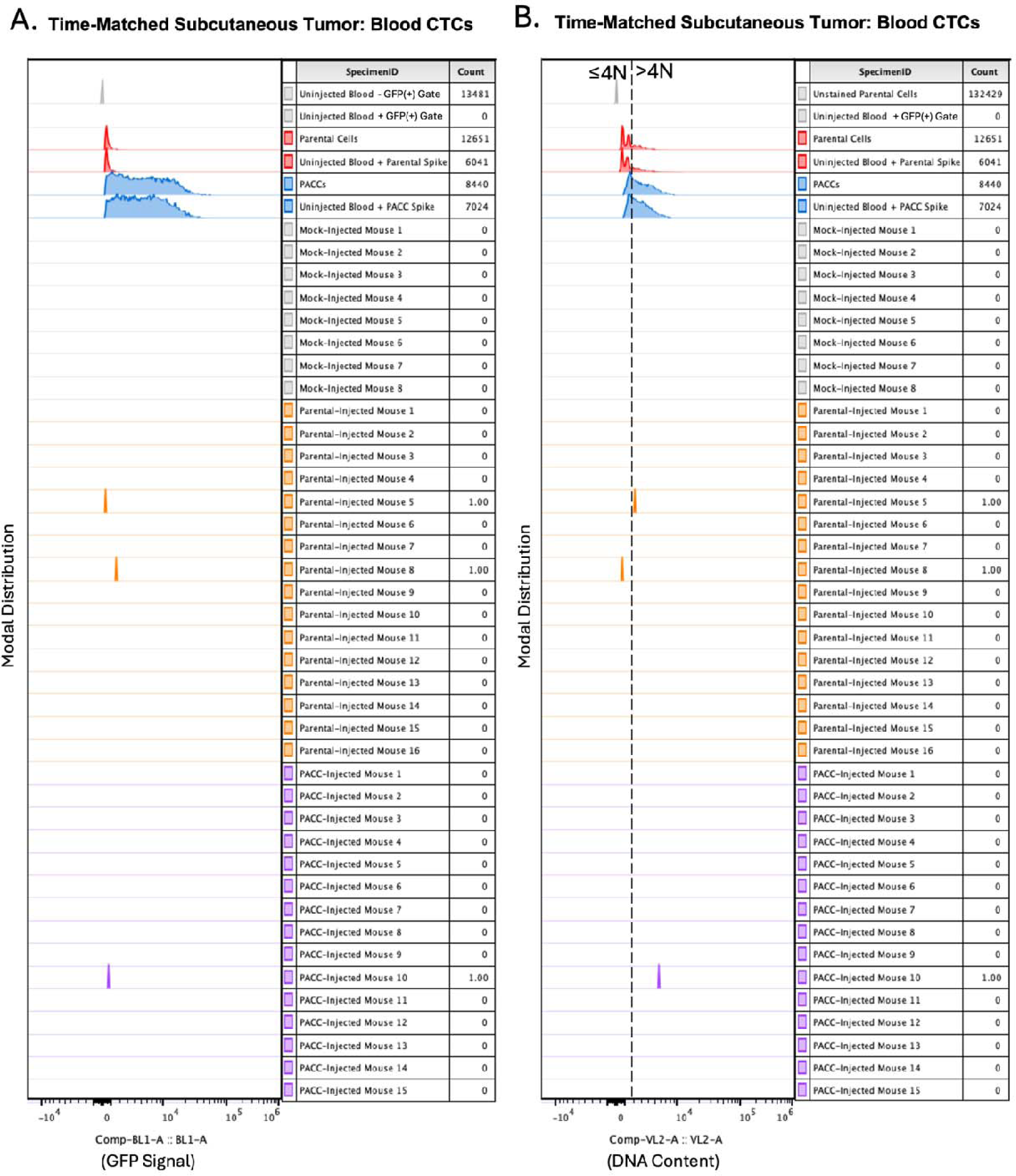
Time-matched, subcutaneous injection of PC3-GFP-Luc parental population vs. PACC-enriched population: A) Raw cytometric data reporting GFP+ signal across all blood samples. B) Raw cytometric data reporting DNA content of GFP+ cells across all blood samples.

**Supplemental Figure S2:**
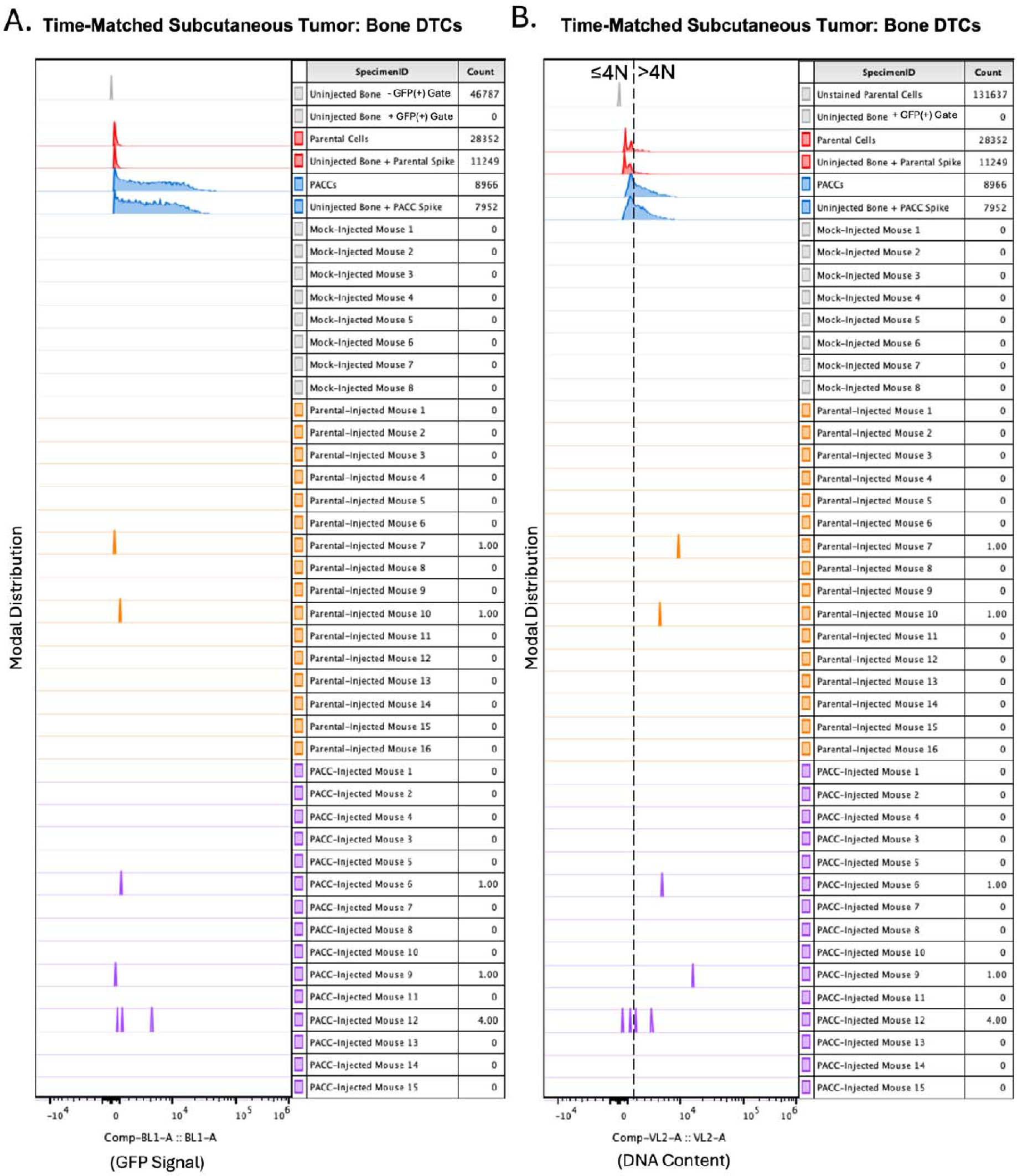
Time-matched, subcutaneous injection of PC3-GFP-Luc parental population vs. PACC-enriched population: A) Raw cytometric data reporting GFP+ signal across all bone marrow samples. B) Raw cytometric data reporting DNA content of GFP+ cells across all bone marrow samples.

**Supplemental Figure S3:**
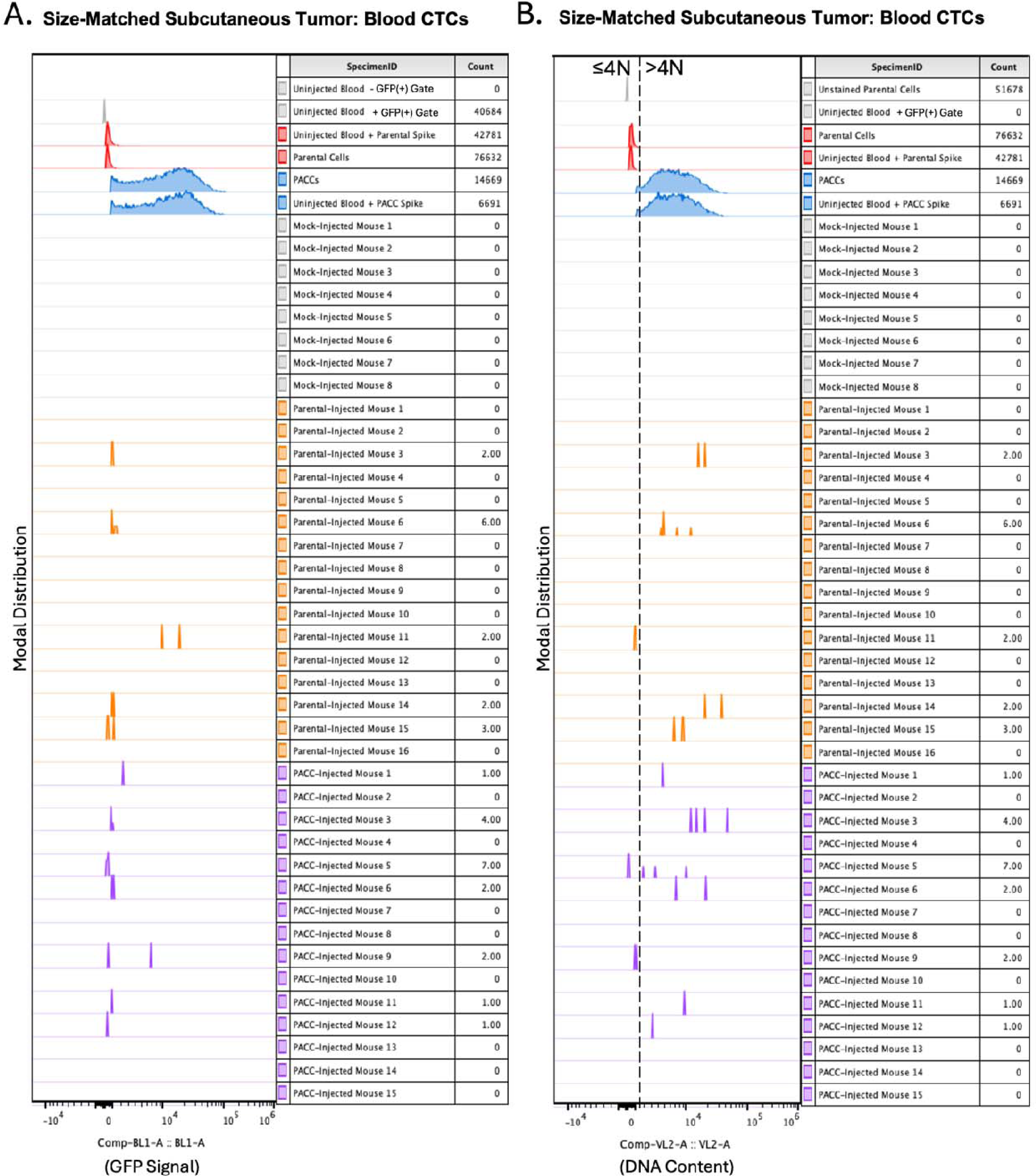
Size-matched, subcutaneous injection of PC3-GFP-Luc parental population vs. PACC-enriched population: A) Raw cytometric data reporting GFP+ signal across all blood samples. B) Raw cytometric data reporting DNA content of GFP+ cells across all blood samples.

**Supplemental Figure S4:**
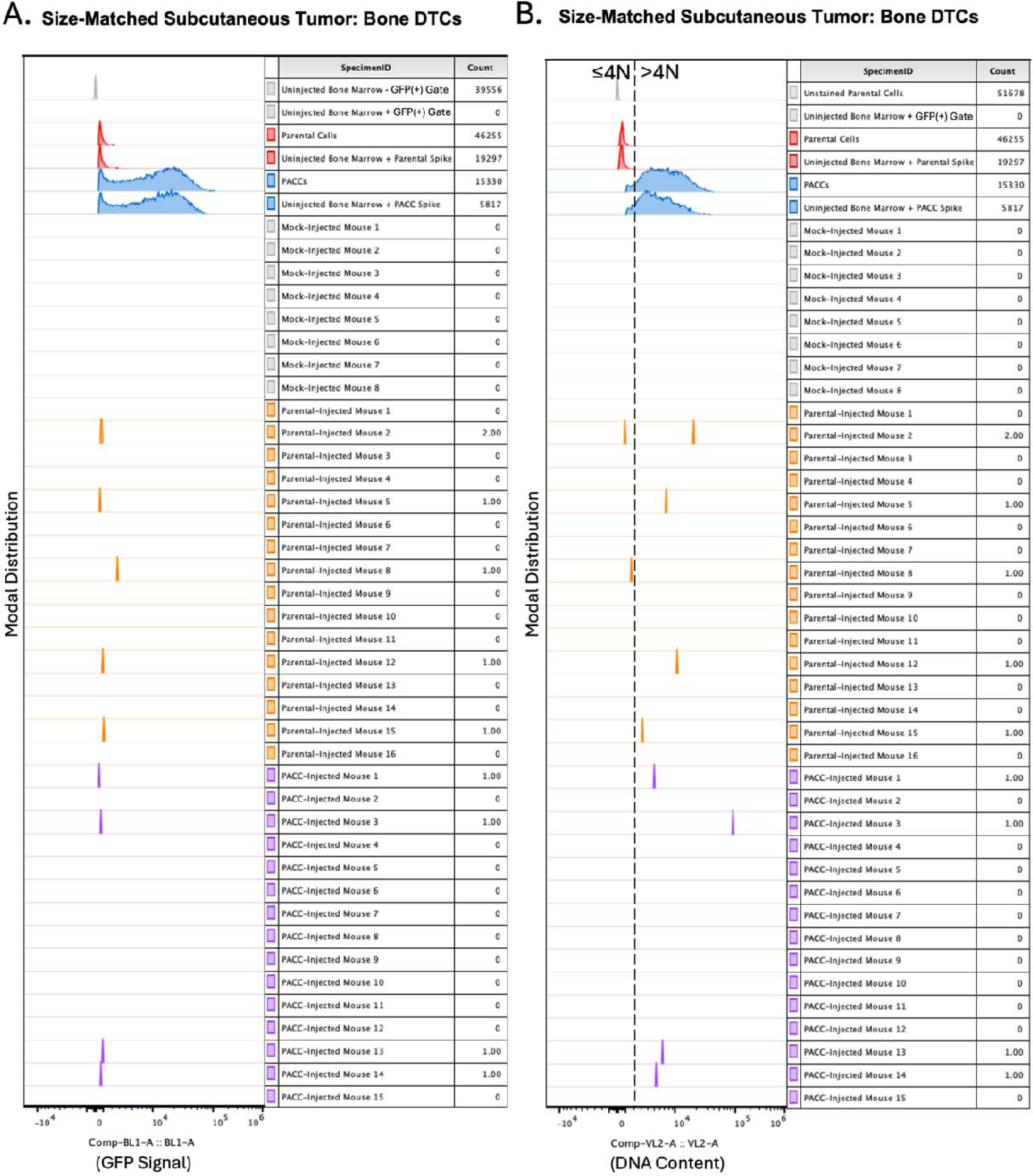
Size-matched, subcutaneous injection of PC3-GFP-Luc parental population vs. PACC-enriched population: A) Raw cytometric data reporting GFP+ signal across all bone marrow samples. B) Raw cytometric data reporting DNA content of GFP+ cells across all bone marrow samples.

**Supplemental Figure S5.**
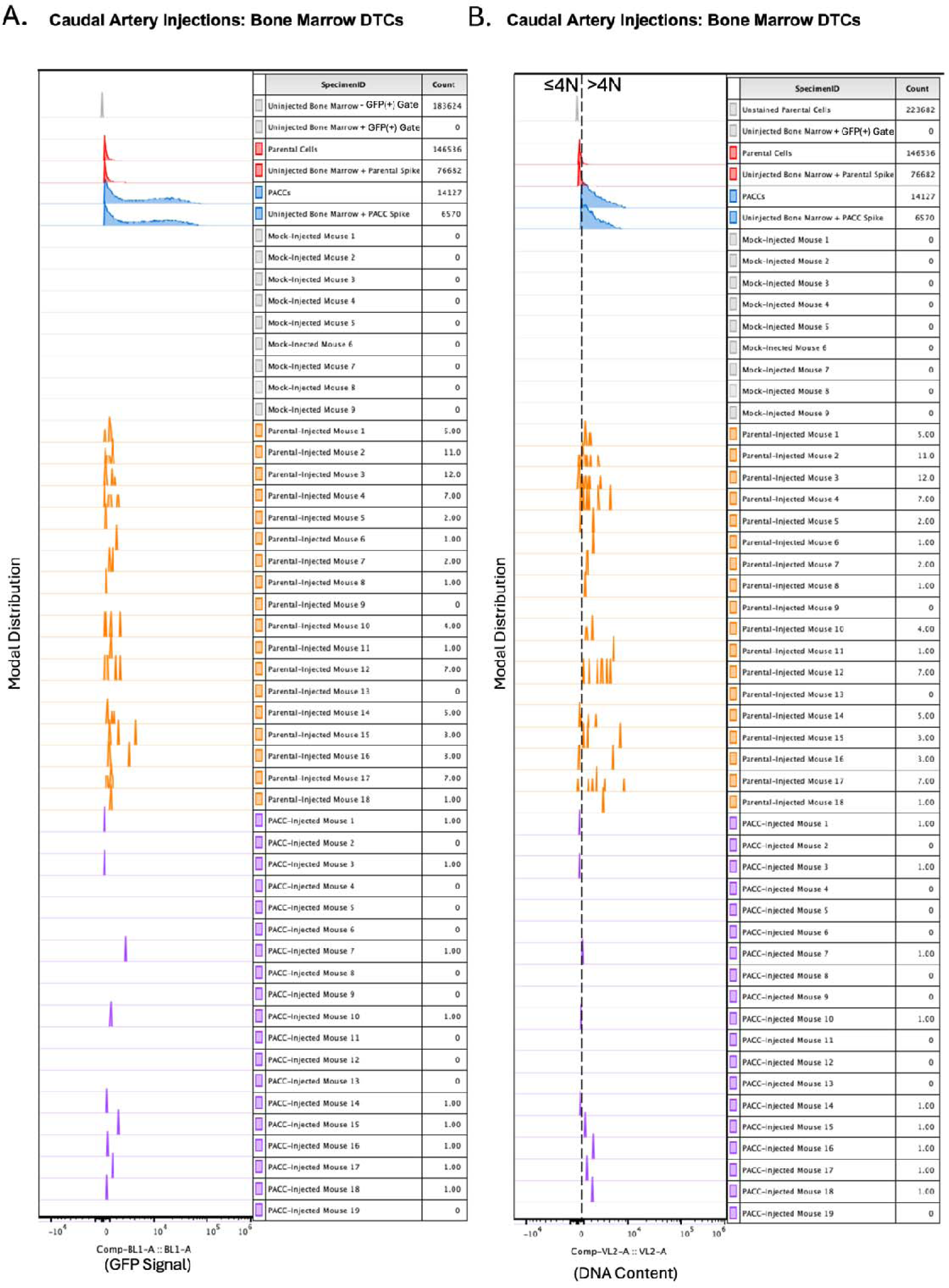
Caudal Artery injection of PC3-GFP-Luc parental population vs. PACC-enriched population: A) Raw cytometric data reporting GFP+ signal across all bone marrow samples. B) Raw cytometric data reporting DNA content of GFP+ cells across all bone marrow samples.

**Supplemental Figure S6:**
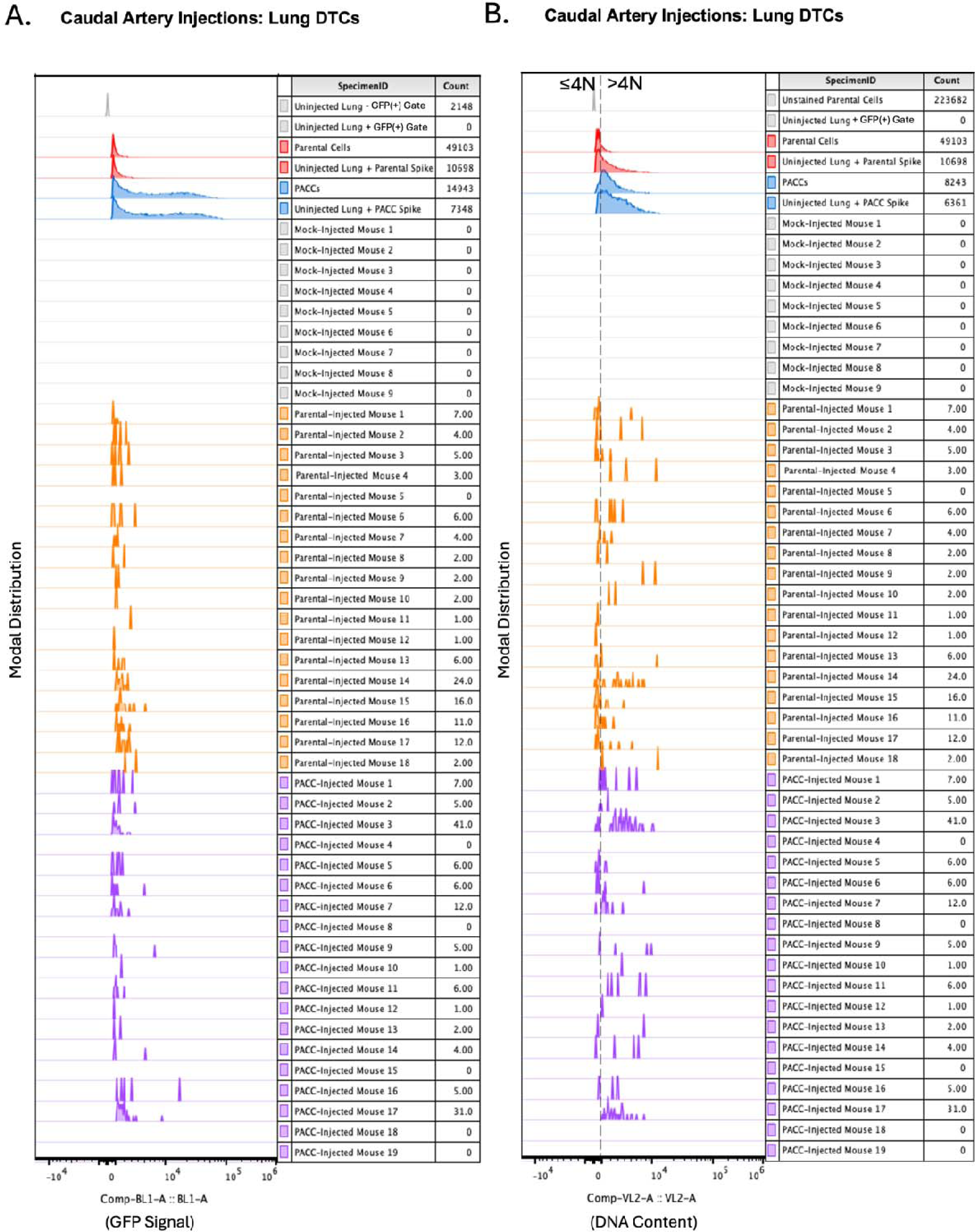
Caudal Artery injection of PC3-GFP-Luc parental population vs. PACC-enriched population: A) Raw cytometric data reporting GFP+ signal across all lung tissue samples. B) Raw cytometric data reporting DNA content of GFP+ cells across all lung tissue samples.

**Supplemental Figure S7:**
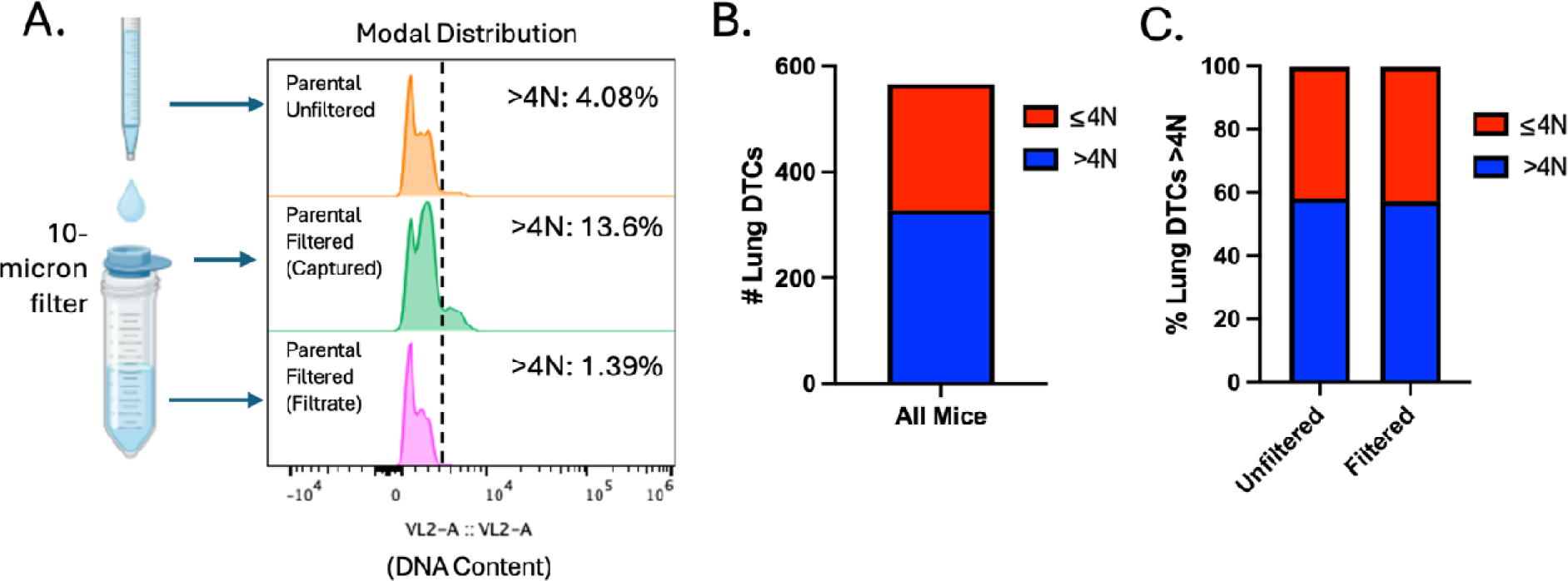
Tail Vein injection of PC3-GFP-Luc parental population vs. PACC-depleted parental population: A) Flow-cytometric ploidy analysis of injected cells per injection group. B) Proportion of lung DTCs with >4N DNA content across all experimental mice. C) Comparative proportions of lung DTCs with >4N DNA content between two injection groups.

**Supplemental Figure S8:**
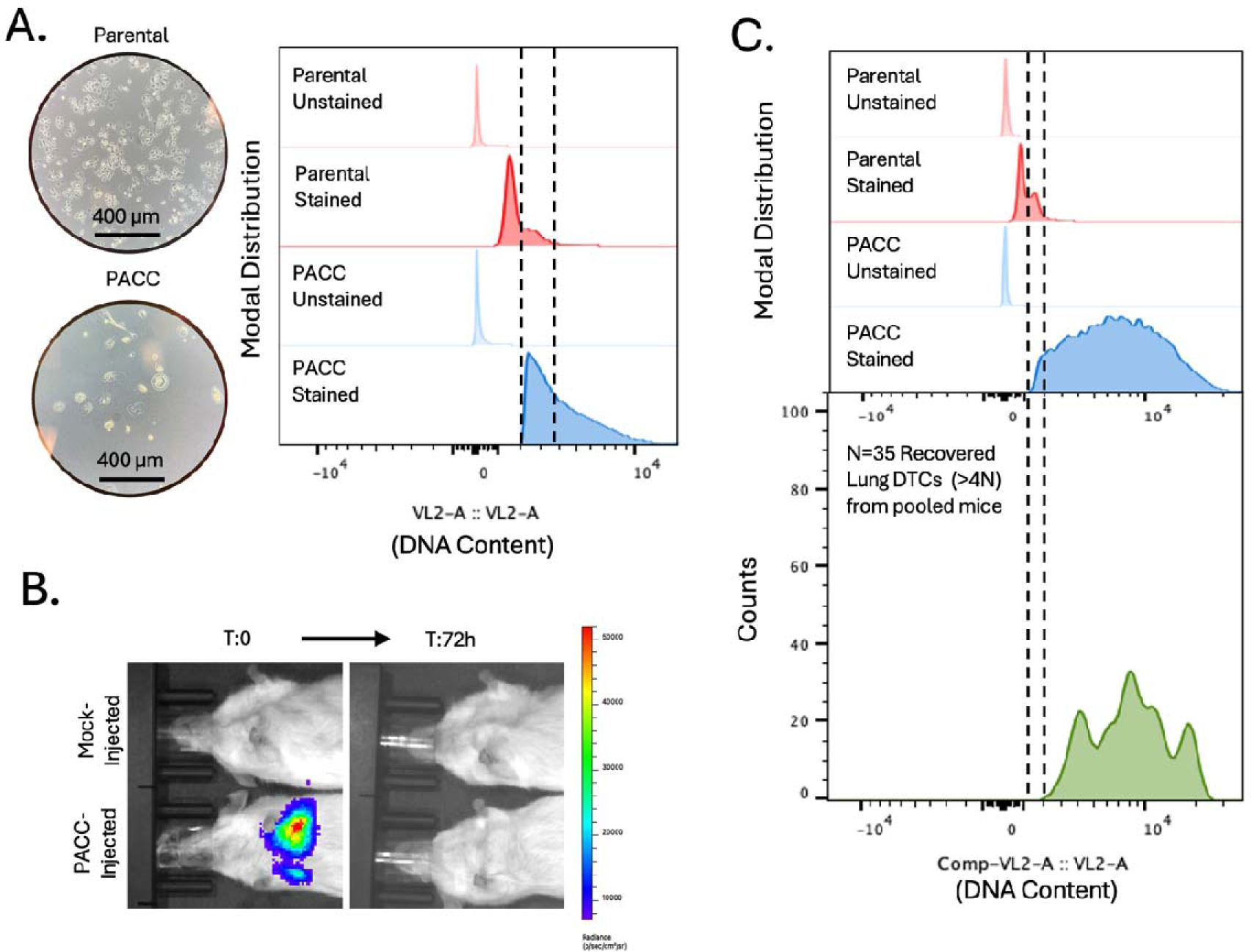
Tail Vein injection of PC3-GFP-Luc parental population vs. PACC-enriched population: A) Light microscopy photos and flow-cytometric ploidy analysis of injected cells per injection group. B) Representative BLI images capturing the cellular distribution and signal intensity immediately following tail vein injection and 3 days following tail vein injection. C) Raw cytometric data reporting the presence of >4N Lung DTCs in a pooled sample 21 days following PACC injection.

**Supplemental Figure S9:**
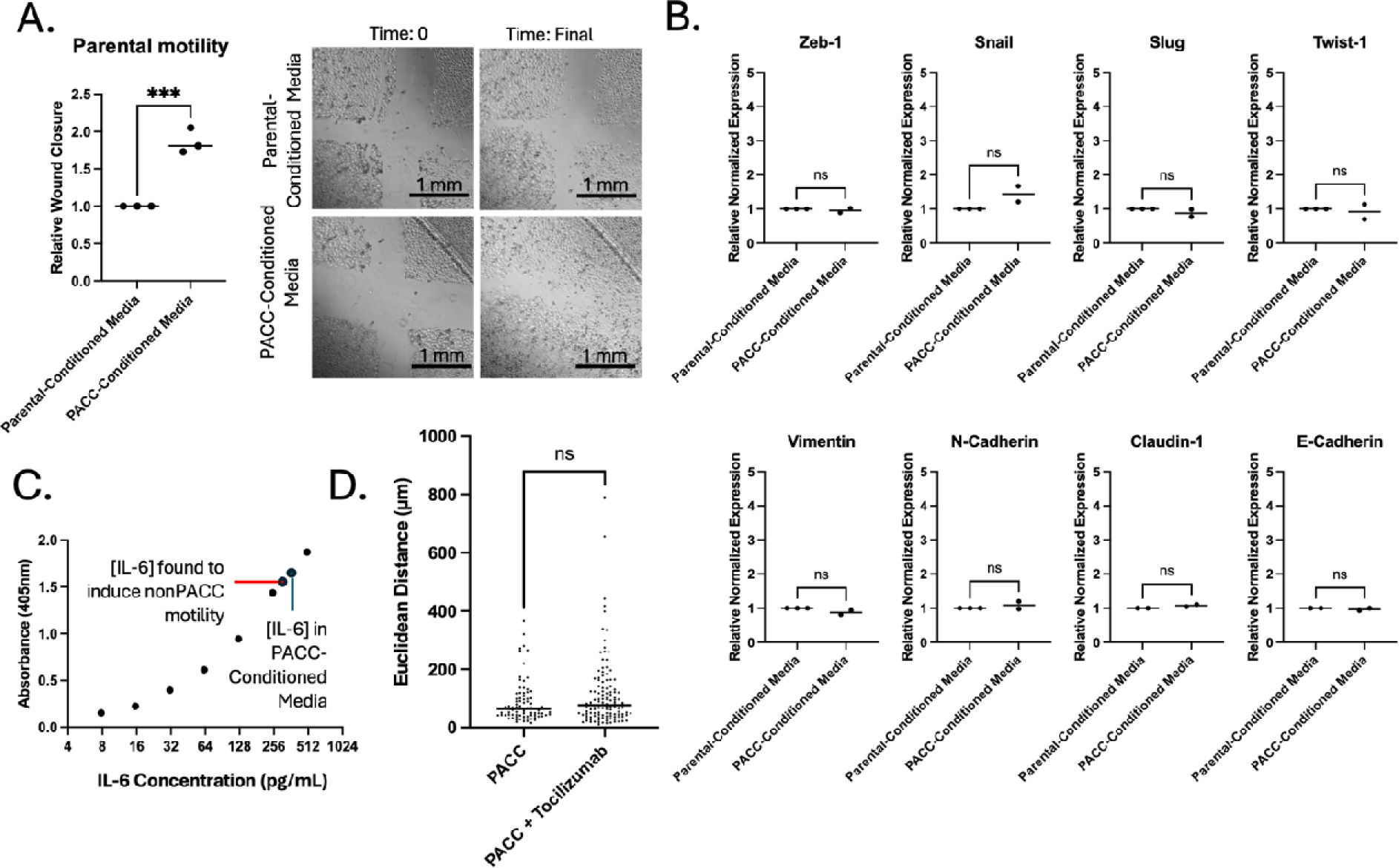
PACCs have a pro-metastatic secretory profile: A) Wound-healing assay to measure the differential motility of a PC3-Luc population exposed to parental-conditioned media vs. PACC-conditioned media, where each reported biological replicate is an average of two technical replicates. B) RNA expression of a panel of EMT markers by RTqPCR in a PC3-Luc parental population exposed to parental-conditioned media vs. PACC-conditioned media, were each biological replicate reported is an average of three technical replicates. C) Elisa assay to measure the concentration of IL6 in PACC conditioned media. D) Single cell tracking to measure the effects of addition of Tocilizumab on the motility of a PACCs.

## Notes

### Competing Interest Statement

M.M.M. has no disclosures. L.T.A.R. has no disclosures. M.J.S has no disclosures. S.P.N. has no disclosures. A. J. Z. has no disclosures. P.K. discloses ownership in Epic Sciences. J.H discloses he is a member of the Clinical Advisory Board of Epic Sciences. K.J.P. discloses that he is a consultant to Cue Biopharma, Inc., an equity holder in PEEL therapeutics, and a founder and equity holder in Keystone Biopharma, Inc. and Kreftect, Inc. S.R.A. discloses that she is an equity holder in Keystone Biopharma, Inc.

